# Healthful choices depend on the latency and rate of information accumulation

**DOI:** 10.1101/465393

**Authors:** Nicolette J. Sullivan, Scott A. Huettel

## Abstract

The drift diffusion model (DDM) provides a parsimonious explanation of decisions across neurobiological, psychological, and behavioral levels of analysis. Although most DDM implementations assume that only a single value guides decisions, choices often involve multiple attributes that could make separable contributions to choice. Here, we fit incentive-compatible dietary choices to a multi-attribute, time-dependent drift diffusion model (mtDDM), in which taste and health could differentially influence the evidence accumulation process. We found that these attributes shaped both the relative value signal and the latency of evidence accumulation in a manner consistent with participants’ idiosyncratic preferences. Moreover, by using a dietary prime, we showed how a healthy choice intervention alters mtDDM parameters that in turn predict prime-dependent choices. Our results reveal that different decision attributes make separable contributions to the strength and timing of evidence accumulation – providing new insights into the construction of interventions to alter the processes of choice.

## Main

Simple choices, like those between food items, have been characterized using sequential integrator models such as the drift (or decision) diffusion model (DDM)^1–4^. In the DDM, choices arise from a process that dynamically integrates evidence for and against each option over time – and a decision is made when the evidence signal reaches the threshold associated with one of the choice options. These models have been enhanced to account for various features of the decision process, allowing them to better explain choices and to generate new insights into cognitive processes. For example, gaze,^5^ and pupil dilation^6^, and neural data^7–11^ have incorporated the influence of attention and neural signals, resulting in improved predictions. See Ratcliff^12^ for a review of advances in the DDM.

A key advantage of these models is their ability to dissociate the influences of distinct cognitive processes, such as distinguishing bias toward one choice option from reductions in the amount of evidence needed before deciding. Although current variants of these models provide highly accurate descriptions of the psychometrics of value-based choices (i.e., describe both choices and their response times in laboratory experiments), they do not account for important potential contributors to the choice process, including distinct contributions of different attributes to a single choice^13,14^.

Here, we present a multi-attribute, time-dependent, drift diffusion model (mtDDM) that modifies the traditional DDM in two ways. First, it estimates the rate of evidence accumulation at each time point (“drift slope”) separately for two attributes, which allows estimation of their unique contributions while controlling for other features of the decision process^15,16^. Second, the mtDDM allows each attribute to begin influencing the decision process at a distinct time (“drift latency”). This builds on previous work in which processing of irrelevant features must be inhibited (e.g., Stroop tasks), potentially through shifts in the drift process or two-stage diffusion processes^17–22^. Similarly, previous efforts to understand the temporal order of events in the brain – such as the timing of automatic and voluntary processes – have enhanced our understanding of cognition and behavior^23,24^.

A potential strength of the mtDDM is its ability to distinguish among different pathways that could each lead to an unhealthy choice. Most commonly, an individual could place a large weight on taste or a small weight on health. Alternatively, and non-exclusively, relatively delayed processing of health information might preclude its consideration in the decision process – leading to unhealthy choices that run counter to the decision-maker’s preferences. There also could be an interaction between decision weights and the timing of processing; e.g., an earlier entry of health information could compensate for a large weighting on the taste attribute. Any of these pathways could result in unhealthy choices, but cannot be differentiated in canonical models.

Dietary choices have several convenient features that make them ideally suited to testing the mtDDM; most importantly, they often involve conflicts between contradictory desires, such as short-term goals related to consumption of a tasty snack and long-term goals related to personal health. In our model, such conflicts can be represented as trade-offs in the separate weights placed on taste and health. Moreover, taste and health have meaningfully different properties, resulting in faster processing of taste than health^13^. This may be because taste is a momentary, immediate, and concrete reward whereas the healthfulness of a food presents only future benefits and involves integration of multiple quantities such as caloric and fat content^25^. Although previous studies have estimated the time at which taste and health are processed^13,14^, they did not disentangle differences in the weight placed on each attribute from differences in timing parameters. By estimating both attributes simultaneously, we can assess their independent contributions.

The predictions of the mtDDM are illustrated in the two plots of Figure 1. Suppose that taste and health enter the decision process at similar times (Fig. 1a). In this case, health influences the value signal toward the healthy option’s boundary early in the decision process, and the healthier option is chosen. In contrast, Figure 1b depicts an identical decision process, except that health’s drift latency is much later, resulting in a large “temporal advantage” such that taste has 300ms longer to influence the value signal. In this example, the value signal has nearly reached the boundary for the tastier option when the health attribute’s latency has been reached. Health therefore would have a more limited time to influence the value signal before a choice is made in favor of the tastier, less healthy option (the top boundary).

**Figure 1.**
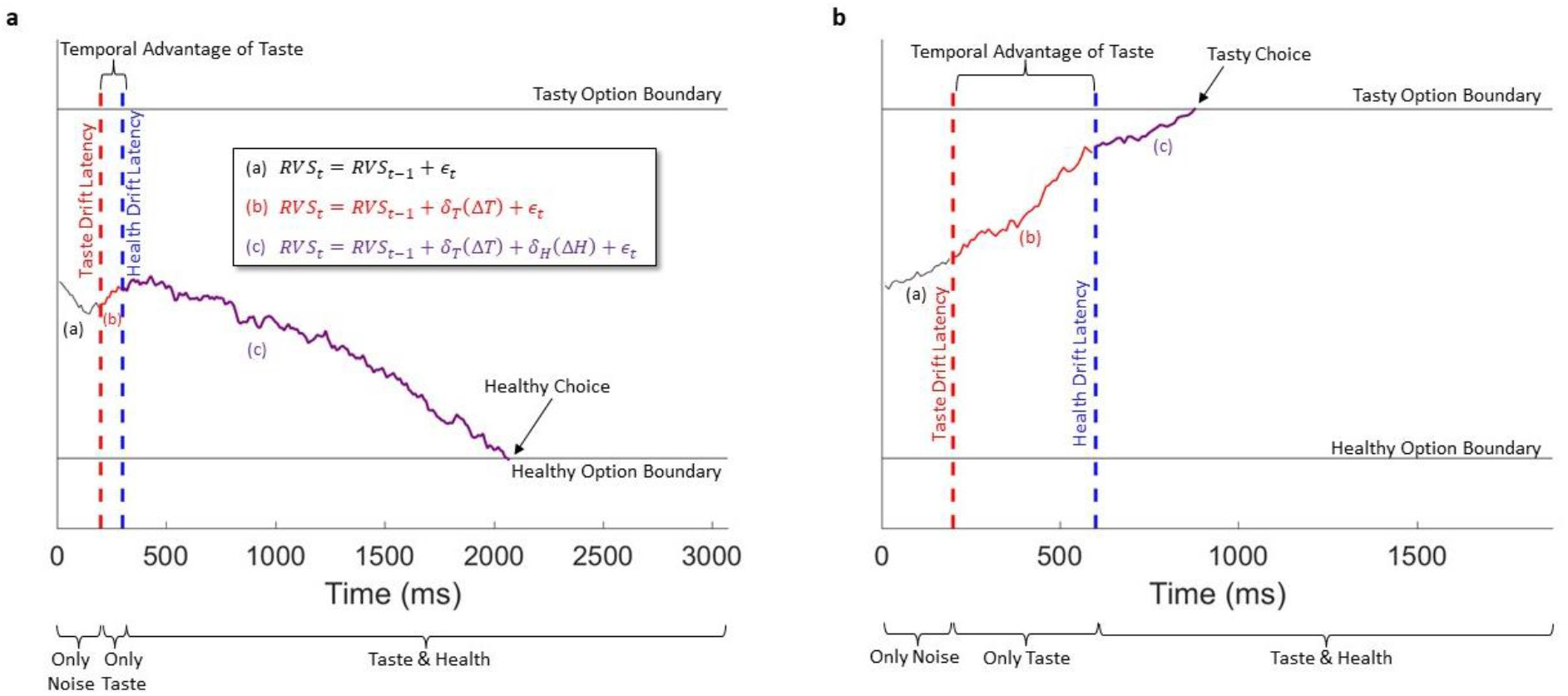
Examples of the decision process modeling within the multi-attribute, time-dependent drift diffusion model (mtDDM). In these example choices between a tasty food (tasty but unhealthy) and a healthy food (healthy but not tasty), a relative value signal (RVS) begins with a value set by a bias parameter (here, zero) and evolves only with noise, *ε*, at every timepoint *t* as depicted in equation and segment “a”. Once the taste latency is reached (red dotted line), the relative (tasty– healthy) taste value, *ΔT*, begins contributing to the RVS at a rate determined by its drift slope *δ*_*T*_ (“b”). After the health attribute latency is reached (“c”), relative health value, *ΔH*, begins contributing to the RVS at a rate determined by its drift slope *δ*_*H*_. At each time point, Gaussian noise *ε* is added to the value signal. A decision is reached when the RVS becomes equal to or greater than the boundary for an item. **(a)** In this example, a simulated RVS path is displayed for a choice in which the difference in taste and health attributes are *ΔT*=1 and *ΔH*=−7, and mtDDM parameters are set to *δ*_*T*_ =0.005 units/ms, *δ*_*H*_=0.0009 units/ms, t*_T_=200 ms, t*_H_=300 ms, bias=0, and tasty option boundary=1 and healthy option boundary=−1. Taste and health enter the decision process at similar times, which leads to an early contribution of the health attribute to the RVS, and a healthy choice. **(b)** In this example, all parameters are the same except that the taste attribute has a large “temporal advantage” of 300 ms. That is, due to a later entry of health to the decision process (at t*_H_=500 ms), it begins contributing to the RVS later than in the previous example. Thus, the tasty option boundary is crossed before the health attribute has a significant influence on choice. Figure adapted from^59^.

We tested the robustness of the mtDDM within an incentive-compatible experiment in which participants made a series of binary choices between two foods that varied on two key attributes: their tastiness and healthfulness (Fig. S1). Two behavioral primes were also employed to shape participants’ dietary goals via attention to either health or taste attributes, respectively. By focusing attention to each attribute in independent participant groups using a between-subjects design, we perturb the decision process, and thus can evaluate how well the mtDDM can adapt to changes in attribute weighting. Because drift slopes have been shown to vary depending on allocation of attention^26^, an intervention that increases attention to one attribute could increase its rate of accumulation and therefore bias choice – independently of any effects of dietary self-control. We hypothesize that increased focus on the primed attribute may also facilitate faster processing of that attribute, and that these speeded latencies could help to facilitate more health-focused choices.

Interventions directed at improving choice have found limited success, especially information-based interventions or those targeted at changing patterns of conscious thought^27,28^. Therefore, it is critical to identify the mechanisms underlying what seem to be failures in dietary self-control – particularly if healthy choices might depend on something other than self-control or preferences^29–32^. For example, if healthy choices are facilitated in part by other features of the decision process, such as the time at which health information is processed, harnessing those features may aid in the development of effective interventions. This paper goes beyond introducing an innovation to sequential integrator models to also suggest ways in which the decision process could be nudged to improve choice.

## Results

### mtDDM Predictions

First, we derived qualitative predictions for how our key new parameters – specifically, the taste and health drift latencies – interact with taste and health drift slopes to influence healthy choices. A series of simulated mtDDMs were performed using an artificial choice set with health and taste values like those in our experimental dataset. Taste’s slope and latency were fixed (to 0.08 units/ms and 500 ms, respectively) and health’s slope and latency was varied so that the influence of changes in the relative (Taste – Health) latency and slope on choice could be visualized (see Supplemental Information for details).

When taste and health drift slopes are equal (Figure 2, purple line) agents made more healthy choices as health latencies became earlier (left to right). This pattern held when taste slopes were larger than health slopes (red lines). Importantly, differences in taste and health drift slope matter less for later health latencies (Figure 2, far left), and matter much more for earlier health latencies (far right). This indicates that as latencies diverge, the influence of slope changes, implying an interaction between the two parameters. Of note, changes in latency had a bigger effect on the attribute whose parameters were fixed, such that an exactly symmetrical effect would be obtained were health parameters to be fixed instead (Fig. S2D). See Supplemental Information for more details.

**Figure 2.**
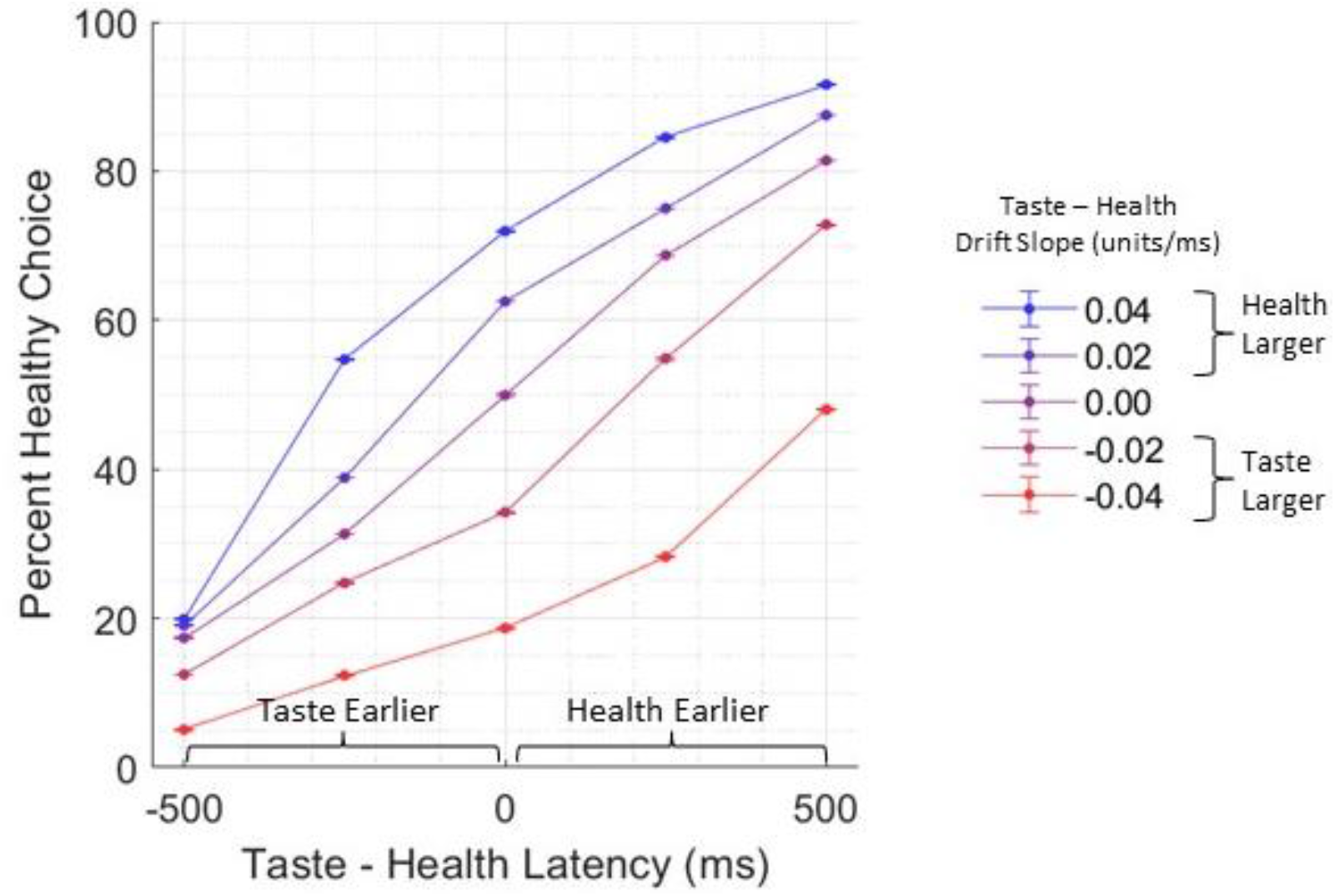
Predicted results of the mtDDM. The proportion of healthy choices (y-axis) predicted by simulations of the mtDDM are plotted for different relative latencies (x-axis). Colored lines show the percent of healthy choices broken out by value difference. Red colors represent simulated agents in which the taste drift slope was larger than health drift slope, and Blue colors represent agents in which the health drift slope was relatively larger.

### Behavioral Results

We performed several tests to ensure that participants were choosing according to their preferences in both prime conditions and that their response times (RTs)fit expected patterns. Choices were significantly related to each option’s reported wanting for both the health-and taste-primed participants (Fig. 3b; mixed effects slope: health prime M=1.02, *d*=1.82, *t*_39_=11.53, *p*<0.001, 95% CI=[0.84 1.19]; taste prime M=1.35, *d*=2.33, *t*_38_=14.57, *p*<0.001, 95% CI=[1.16 1.53]). Logistic regression slopes were statistically significantly smaller in health-primed participants (*d*=− 0.59, *t*_77_=−2.61, *p*=0.01, 95% CI=[−0.59 −0.08]).

**Figure 3.**
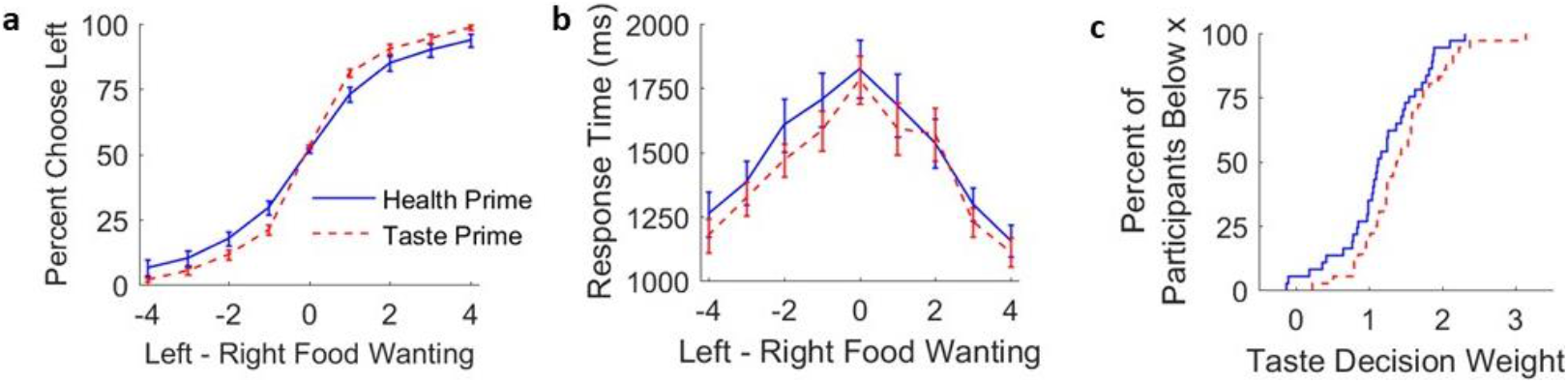
Behavioral results. (**a**) Effects of value difference (Left – Right Food Wanting) on choices. Positive numbers on the x-axis represent cases in which the left item was higher in reported food wanting (mixed effects slope: health prime M=1.02, *d*=1.82, *t*_39_=11.53, *p*<0.001, 95% CI=[0.84 1.19]; taste prime M=1.35, *d*=2.33, *t*_38_=14.57, *p*<0.001, 95% CI=[1.16 1.53]). (**b**) Mean response time (RT) is shown as a function of choice difficulty as measured by the difference between wanting for the right item and for the left item. A difference of zero indicates a difficult choice between two equally-wanted options, and a 4 or −4 indicates an easy choice between items with opposite wanting (mixed effects quadratic slope: health prime M=−45.30, *d*=−1.41, *t*_39_=−8.90, *p*<0.001, 95% CI=[−55.60 −35.01]; taste prime M=−40.34, *d*=−1.86, *t*_38_=−11.62, *p*<0.001, 95% CI=[−47.37 −33.31]). (**c**) Cumulative distribution function illustrating taste decision weights by prime condition broken out by whether the participant was primed for health (blue) or taste (red) goals (means 1.18 vs. 0.13, *d*=2.24, *t*_77_=9.94, *p*<0.001, 95% CI=[0.84 1.27]). For plots A and B, error bars represent standard error of the mean. For all plots, the health prime condition is represented by the blue solid line and taste prime condition by the red dotted line.

Faster RTs for conflict than non-conflict trials (Fig. S3a; M=1557.59 ms, 1635.57 ms; paired t-test of log(RTs) d=−0.17, t(77)=−3.34, p=0.001, 95% CI=[−0.08 −0.02]) were driven by fast unhealthy choices, as healthy choice RTs were markedly longer (Fig. S3b; M=1917 ms, 1493 ms; paired t-test of log(RTs) d=0.61, t(76)=7.33, p<0.001, 95% CI=[0.15 0.27]). This is expected from any DDM with separate weights on taste and health and is the result of the accumulating advantage of taste information during the decision process (see “Response Times by Choice and Trial Type” section of Supplement). RTs increased with choice difficulty – as measured by a difference of zero in reported wanting indicating a difficult choice between two equally-wanted options – for both groups (Fig. 3c; mixed effects quadratic slope: health prime M=−45.30, *d*=−1.41, *t*_39_=−8.90, *p*<0.001, 95% CI=[−55.60 −35.01]; taste prime M=−40.34, *d*=−1.86, *t*_38_=−11.62, *p*<0.001, 95% CI=[−47.37 −33.31]). Quadratic regression slopes were not statistically significantly different between the taste than health prime (*d*=−0.18, *t*_77_=−0.80, *p*=0.43, 95% CI=[−17.29 7.36]), nor were average RTs (means 1628 vs. 1554 ms, *d*=0.13, *t*_77_=0.59, *p*=0.56, 95% CI=[−175.39 323.30]). These results indicate that individuals used value to guide choice in both conditions, and that health-primed participants weighted wanting less than taste-primed participants.

We next estimated the influence of our behavioral prime on choice using each food’s taste and health differences, to estimate the weight each participant placed on taste and health information in their decisions, and how this changed depending on the prime they received. We found that health-primed participants placed significantly less weight on taste information (Fig. 3d; means 1.18 vs. 0.13, *d*=2.24, *t*_77_=9.94, *p*<0.001, 95% CI=[0.84 1.27]), which resulted in a marginal increase in the proportion of healthy choices in the health prime condition, as assessed by comparing their log-transformed values (means=0.26, 0.18; d=0.44, t(75)=1.94, p=0.057, 95% CI=[−0.01 0.83]).

### Fitted parameters of the mtDDM

Using participants’ choices and RTs, we fit five mtDDM parameters (Table 1; see Fig. S4 for parameter distributions). These parameters were the weight placed on taste and health during option comparison (“Drift Slope”, δ_T_ and δ_H_), the time required for taste and health to enter option comparison (“Drift Latency”, t*_T_ and t*_H_), and the evidence required to make a choice (“Boundary”, b). Taste drift slopes were larger than health drift slopes (*d*=1.83, t_78_=10.91, *p*<0.001, 95% CI=[0.04 0.05]), reflecting a greater emphasis on taste information in evidence accumulation. Together, these results confirm findings from the previous literature that taste has an overall “weighting advantage” in the choice process. Drift latency was significantly earlier for taste than for health (*d*=−0.98, t_78_=−6.22, *p*<0.001, 95% CI=[−603 −311]). These results indicated that taste information has two advantages during the decision process – both an earlier entry and a greater contribution to evidence accumulation – that together may explain taste’s greater influence on the relative value signal and subsequent decisions.

**Table 1.** Average best-fitting mtDDM parameters.

| Parameter | Mean (SD) | Min | Max |
| --- | --- | --- | --- |
| Taste Drift Slope, $\delta_T$ (units/ms) | 0.062 (0.030) | 0.003 | 0.148 |
| Health Drift Slope, $\delta_H$ (units/ms) | 0.017 (0.018) | 0 | 0.086 |
| Taste Drift Latency, $t^*_T$ (ms) | 409 (245) | 10 | 1920 |
| Health Drift Latency, $t^*_H$ (ms) | 866 (611) | 20 | 2750 |
| Boundary, $b$ (units) | 1.414 (0.200) | 1.088 | 1.950 |

### Correlation between parameters

There was not a statistically significant correlation between taste and health drift slopes (Pearson *ρ*=−0.12, *p*=0.29) nor between taste and health drift latencies (*ρ*=0.02, *p*=0.87). Each attribute’s drift slope and latency were not statistically significantly correlated (Taste, Pearson *ρ*=0.11, *p*=0.32; Health, Pearson *ρ*=−0.11, *p*=0.32). Boundary width, typically linked to response caution^33–37^(although see^38^) was not statistically significantly related to health drift slopes or taste drift latencies (*p*>0.12). However, larger boundary width was marginally correlated with smaller taste drift slopes and with later health drift latencies (*δ*_*T*_, Pearson *ρ*=−0.20, *p*=0.07; *t*\*_*H*_, Pearson *ρ*=0.35, *p*=0.001). This could arise from artifactual interdependencies between model parameters^39–41^ resulting in trade-offs between drift rates, latencies, and boundaries. However, we did not find a statistically significant correlation between slope and boundary parameters in in the recovery dataset (Taste Slope and Boundary, Pearson correlation ρ=−0.09, p=0.36; Health Slope and Boundary, Pearson ρ=−0.17, p=0.09), but did for latency and boundary (Taste Latency and Boundary, Pearson correlation ρ=0.37, p<0.001; Health Slope and Boundary, Pearson ρ=0.25, p=0.01) Alternatively, individuals who process health information later – or who had a smaller contribution of taste information during evidence accumulation – may have required more evidence to make a choice. This may allow some individuals to compensate for a late health drift latency and still make a healthy choice – a hypothesis we investigate in the penultimate Results section. See Fig. S5 for parameter correlations.

### Model validations and comparisons

To validate the model, a parameter recovery was performed (see Supplemental Results and Fig. S6). This ensured that a large drift slope for one attribute would not lead to inaccurately fast estimate of that attribute’s latency. The mtDDM also was tested against and performed better, as assessed by lower Bayesian Information Criterion (BIC), than four alternative models (see Table S1 and Supplemental Results): one in which only slopes, and not latencies varied by attribute (multi-attribute DDM, mDDM); one in which only latencies, and not slopes, varied by attribute (latency DDM, latDDM); one in which attributes vary in the times when they stop rather than start contributing to the value signal (stopping time DDM, stDDM); and one in which taste and health have equal slopes and latencies (simple DDM, sDDM). Lastly, we tested, but do not find evidence for, the possibility that latencies differences could arise from differences in unhealthy and healthy choice non-decision times (see Supplemental Results).

#### mtDDM vs. single-latency model

We next test the ability of the mtDDM to explain choices and RTs better than a model without separate latencies. This DDM was identical to the mtDDM, but that additionally assumed that taste and health enter the decision process simultaneously. This is functionally equivalent to a simple DDM with one relative value signal (taste + health; see Supplemental Methods)

To compare models, we obtained a BIC using both choices and RTs to classify a correct prediction. Critically, the BIC penalizes the mtDDM for having two latency parameters. The mtDDM performs better (mean BIC values 1111 vs. 1143; d=−0.33, t(78)=−7.95, p<.001, 95% CI=[−40 −24]) for 95% (75 of 79) participants, indicating that the addition of attribute-wise latency parameters generates an improvement in model performance, capturing variance in choices and RTs that a single-latency model cannot.

#### mtDDM parameters are proportional to their influences on choice

To validate that the mtDDM accurately reflects participants’ choices, we next tested the relationship between drift slopes and the weight placed on each attribute in choice; this relationship is expected because an attribute’s drift slope represents the weight placed on that attribute throughout the choice process. These analyses were performed using cross-validated estimation, in which mtDDM parameters were fit using one half of a participant’s data, and were used to predict choice in the other half of data (see Supplemental Methods). First, we estimated the relationship between taste and health drift slopes and their decision weights using a linear regression and found that a participant’s relative drift slope (taste – health) fitted to one half of trials was correlated with the relative weight (taste – health) during choice in the other half of trials (Fig. 4a; taste – health weights; R^2^=0.67, slope=19.57, *p*<0.001). Furthermore, an increased likelihood of healthy choices in Conflict Trials (when one food was healthier, but less tasty, than the other) was related to relatively smaller taste and larger health drift slopes (Fig. S7a; *R*^*2*^=0.67, slope=−5.14, *p*<0.001). Together, these results confirmed that drift slopes reflected the weight participants placed on taste and health during choice and captured a large proportion of variance in healthy choices, suggesting a correct fitting of the model to choices.

**Figure 4.**
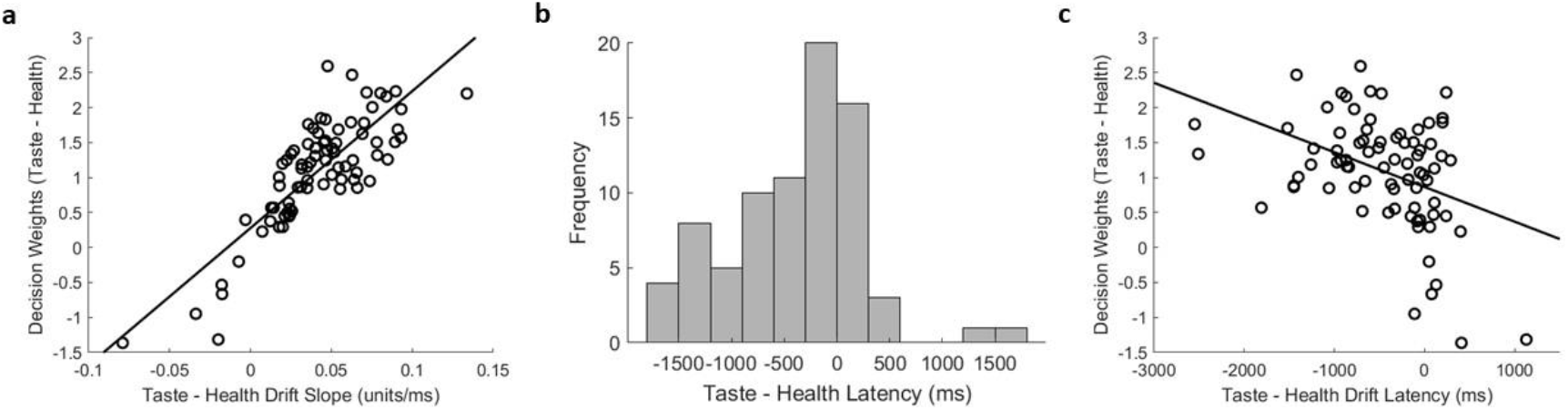
Attribute drift latency parameters related to choice. (**a**) Histogram depicting the distributions of best-fitting taste and health drift slopes (δ_T_ and δ_H_) across participants. (**b**) Taste and health drift latency shown as a function of their logistic decision weights. Lines represent best-fitting linear regression lines. (**c**) Relative (taste – health) drift latency shown as a function of proportion of healthy choices. Lines represent best-fitting linear regression lines.

Because the an attribute entering the decision process earlier influences the relative value signal for longer, it should have a greater influence on choice, all else being equal (see Figures 1 and 2). However, latency differences could fail to drive choices if those differences were very small compared to the overall choice period or if the drift slopes were so large that they dominated the choice process. These concerns are partially addressed by the mtDDM’s better fit compared to the mDDM, and by noting that the taste information enters the choice process approximately 450 ms earlier than health information (Fig. 4b; one-sample t-test vs. 0, d=−0.70, t(78)=−6.22, p<0.001, 95% CI=[−603 −311]). This result, combined the mtDDM’s lower BICs and the successful parameter recovery, provides converging evidence that drift latencies themselves do differ by attribute, and that taste has a temporal advantage in the decision process.

We next confirmed that, across participants, taste’s temporal advantage was related to an increased decision weight on health, relative to taste – again, using cross-validation fitted DDM parameters (Fig. 4c; *R*^*2*^=0.16, slope=5×10^−4^, *p*<0.001) and thus more healthy choices (Fig. S7b; *R^2=^*0.18, slope=1×10^−4^, *p*<0.001). A robust regression approach to confirmed that this relationship held even when excluding outliers in Figure 4c (slope=6×10^−5^, *p*=0.02).

These results indicate that the influence of taste and health on choice depends on the time at which each attribute began to influence the decision process. They also provide an additional explanation for apparent failures of dietary self-control: for many individuals, health information enters the decision process too late (relative to taste information) to drive choices toward the healthier option.

### Drift slope and attribute latency have independent influences on healthy choice

We had hypothesized that drift slope and latency exert independent influences on choice, even when controlling for each other. To test this, we estimated a series of multiple linear regressions using drift slope and latency differences (Taste – Health) to predict individual differences in the proportion of healthy choices made (Table 2). To control for response caution, boundary width was also included. This method tests whether drift slopes and latencies explain different types of variance in the proportion of healthy choices participants made. This analysis was performed using drift slopes fitted using the mDDM, latencies fitted using the latDDM, and boundary width fitted using the sDDM. As expected, drift slopes and latencies predicted individual differences in proportion of healthy choices in the full model. All variables together explained a much larger proportion of the variance in healthy choices than any other model (72%; Model 5 in Table 2); of note, we then performed the same prediction using mDDM drift slopes (i.e., the single-latency model), which explained less variance in healthy choices a model that included latency differences as well (57%; see Model 1 vs. 5 in Table 2).

**Table 2.**
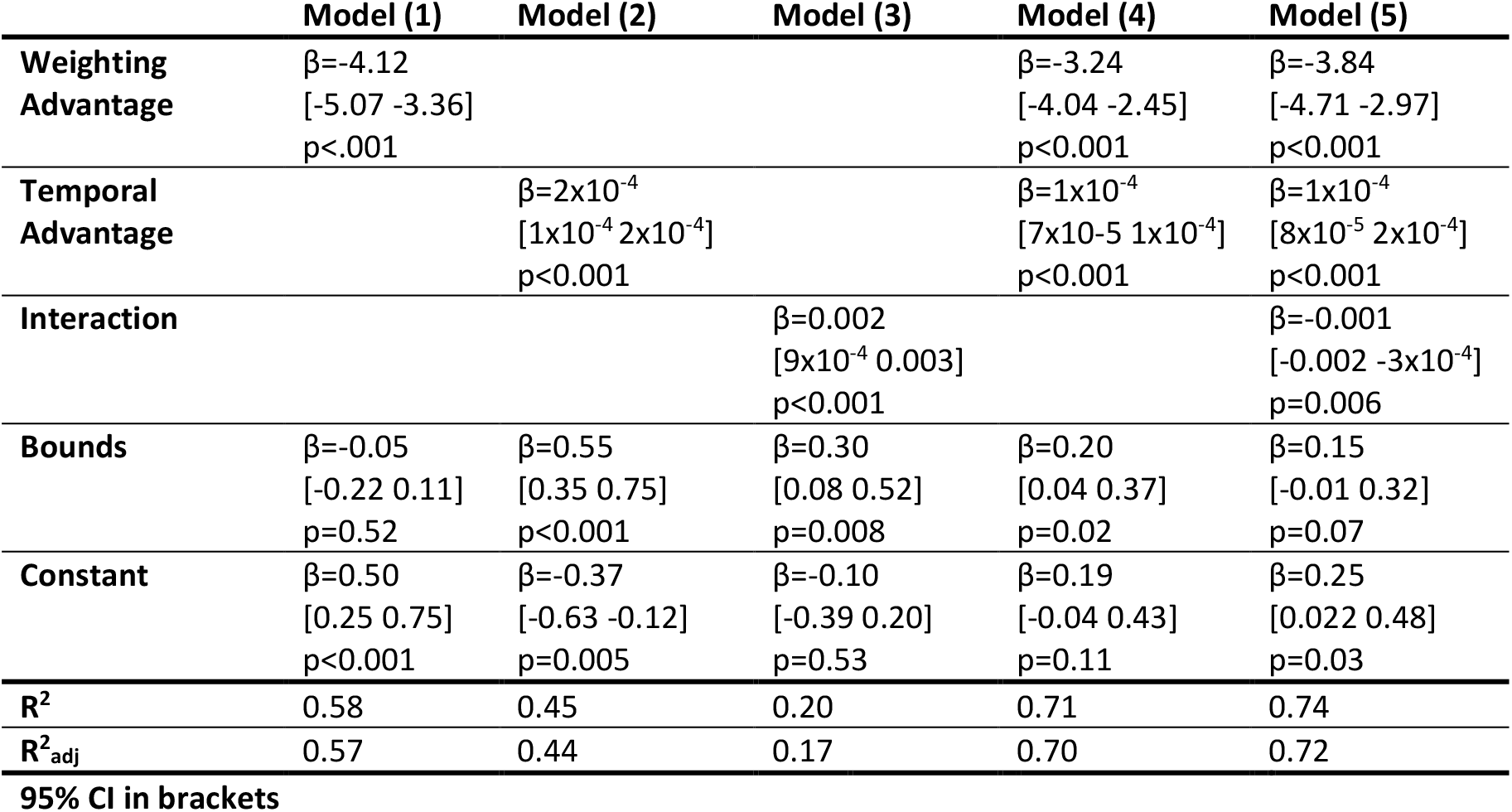
Relationship between proportion healthy choices and fitted mtDDM parameters Weighting advantage was fit using mDDM slopes, Temporal advantage was fit with latDDM latencies, and Bounds were fit with sDDM bounds.

|  | Model (1) | Model (2) | Model (3) | Model (4) | Model (5) |
| --- | --- | --- | --- | --- | --- |
| **Weighting Advantage** | $\beta=-4.12$<br>[-5.07 -3.36]<br>$p<0.001$ | | | $\beta=-3.24$<br>[-4.04 -2.45]<br>$p<0.001$ | $\beta=-3.84$<br>[-4.71 -2.97]<br>$p<0.001$ |
| **Temporal Advantage** | | $\beta=2 \times 10^{-4}$<br>[$1 \times 10^{-4}$ $2 \times 10^{-4}$]<br>$p<0.001$ | | $\beta=1 \times 10^{-4}$<br>[$7 \times 10^{-5}$ $1 \times 10^{-4}$]<br>$p<0.001$ | $\beta=1 \times 10^{-4}$<br>[$8 \times 10^{-5}$ $2 \times 10^{-4}$]<br>$p<0.001$ |
| **Interaction** | | | $\beta=0.002$<br>[$9 \times 10^{-4}$ 0.003]<br>$p<0.001$ | | $\beta=-0.001$<br>[-0.002 -$3 \times 10^{-4}$]<br>$p=0.006$ |
| **Bounds** | $\beta=-0.05$<br>[-0.22 0.11]<br>$p=0.52$ | $\beta=0.55$<br>[0.35 0.75]<br>$p<0.001$ | $\beta=0.30$<br>[0.08 0.52]<br>$p=0.008$ | $\beta=0.20$<br>[0.04 0.37]<br>$p=0.02$ | $\beta=0.15$<br>[-0.01 0.32]<br>$p=0.07$ |
| **Constant** | $\beta=0.50$<br>[0.25 0.75]<br>$p<0.001$ | $\beta=-0.37$<br>[-0.63 -0.12]<br>$p=0.005$ | $\beta=-0.10$<br>[-0.39 0.20]<br>$p=0.53$ | $\beta=0.19$<br>[-0.04 0.43]<br>$p=0.11$ | $\beta=0.25$<br>[0.022 0.48]<br>$p=0.03$ |
| **R<sup>2</sup>** | 0.58 | 0.45 | 0.20 | 0.71 | 0.74 |
| **R<sup>2</sup><sub>adj</sub>** | 0.57 | 0.44 | 0.17 | 0.70 | 0.72 |
| **95% CI in brackets** |  |  |  |  |  |

Next, we used a stepwise linear regression to test which DDM parameters result in the best-fitting model. All DDM parameters from the above model (taste and health slope and latency differences, as well as the sDDM’s temperature parameter and bounds) were added to the regression to predict an individual’s proportion of healthy choices. The best-fitting resulting model was one that included slope difference, latency difference, and their interaction (R^2^_adj_=0.71, F(1,74)=64.80, p<0.001). This further indicates that both slope and latency provide independent contributions to explaining healthy choice.

We next performed a bootstrap mediation analysis^42^ to estimate the additional contribution of drift latency to the proportion of healthy choices. Health drift latency significantly reduced health drift slope’s influence on healthy choice decision weights by 8% (s.e.=.12, p<0.001, 95% CI=[7.66 8.13]). Taste drift slope’s prediction of healthy choices was improved, not reduced, by the inclusion of taste latency (−13% s.e.=0.45, p<0.001, 95% CI = [−14.30 −12.52]). Collectively, these results indicate that individual differences in healthy dietary choice was related to both the drift slope and latency parameters of the mtDDM when controlling for the effects of each other, reflecting their independent contributions.

### Longer RTs associated with greater influence of health

The above findings suggest that longer RTs could increase the likelihood of a healthy choice, as they would allow slower-processed values like health more time to influence the value signal. To test this, we estimated the relationship between individual trial RT and healthy choice in Conflict Trials. We found that longer RTs were associated with an increased likelihood of selecting the healthier food (Table S2, Model 1; mixed-effects logistic regression *R*^*2*^_*adj*_=0.37, log(RT) slope=0.61 (s.e.=0.05), *t*_11698_=11.85, *p*<0.001), which holds when controlling for the reported wanting of the healthy, relative to tasty, option (Table S2, Model 2; *p*<0.001). To assess whether this relationship held across participants, we estimated this regression using average log-transformed conflict trial RTs to predict the proportion of healthy choices made and found the same relationship (robust regression slope=0.09, *t*_76_=2.01, *p*=0.048). This regression was not significant when using non-conflict trial RTs to predict the proportion of healthy choices in Conflict Trials (slope=0.03, *t*_76_=0.56, *p*=0.58). This indicates that longer RTs were correlated with increased likelihood of healthy choices both within and across participants.

Next, we assessed whether this varies by individual mtDDM parameters. If longer RTs promoted healthy choices because they allowed slower-processed health information longer to influence the decision process, then individuals with earlier health latencies would have been less influenced by longer RTs. To investigate this, we first added cross-validation fitted health drift latencies to the previous model predicting healthy choice by RT. RTs remained a significant predictor of healthy choice (Table S2, Model 3; mixed-effects logistic regression, *R*^*2*^_*adj*_=0.48, log(RT) slope=0.39 (s.e.=0.05), *t*_5846_=5.21, *p*<0.001; wanting slope=0.92 (s.e.=0.07), *t*_5846_=12.60, *p*<0.001; t*_H_ slope=0.006 (s.e.=0.002), *t*_5846_=−2.76, *p*=0.006). This indicates that after controlling for the weight of health and taste, RTs continued to explain additional variance in healthy choice.

To assess the interplay between latency and RT, we added interaction terms for RTs and health drift latency. If slower health drift latencies require longer RTs to increase the likelihood of a healthy choice, we would see an interaction between health drift latency and RT. We indeed find that the influence of drift latencies on healthy choice depended on a trial’s RT; longer RTs were associated with increased likelihood of a healthy choices with late health drift latencies. Further, RT’s predictive power was reduced by a third and was no longer significant when drift latency-RT interactions were included in the regression (Table S2, Model 4; mixed-effects logistic regression, *R*^*2*^_*adj*_=0.46, log(RT) slope=0.13 (s.e.=0.12), *t*_5846_=1.05, *p*=0.29; wanting slope=0.92 (s.e.=0.07), *t*_5846_=12.52, *p*<0.001; t*_H_ slope=−0.03 (s.e.=0.010), *t*_5846_=−3.33, *p*<0.001; log(RT) x t*_H_ slope=0.003 (s.e.=0.001), *t*_5846_=−2.77, *p*=0.006). The inclusion of this interaction term resulted in a statistically significant reduction in the influence of RT on healthy choice, as assessed by 1,000 iterations of bootstrap mediation analysis^42^ (mean=30% path strength reduction (s.e.=0.17%) *p*<0.001 95% CI=[29.84% 29.17%]). These results indicate that longer RTs may promote healthful choices by allowing slower-latency health information to contribute to the value accumulation process.

### Dietary primes alter evidence accumulation

Finally, we examined the effects of our two dietary primes – a taste prime and a health prime – on the decision process. There were no statistically significant log(RT) differences between prime groups in Conflict Trials (*p*=.81), Non-Conflict Trials (*p*=.94), or for healthy or unhealthy choices (*p*>.55). Taste drift slopes were smaller for health- than taste-primed participants (Fig. S8a; means 0.06 vs. 0.07, *d*=−0.48, *t*_77_=−2.12, *p*=0.04). Log-transformed taste drift slopes were also relatively smaller than health drift slope for health primed participants (δ_T_-δ_H_; Fig. S8b; means 0.04 vs. 0.05, *d*=−0.47, *t*_77_=−2.08, *p*=0.04). No other parameter differed statistically significantly between condition (p > 0.27). Together, these results provide no credible evidence that the health prime slowed the overall decision process, per se, but instead that the prime influenced the degree to which taste information influenced the value signal, both in absolute terms and relative to health information.

## Discussion

Sequential integrator models such as the DDM have been used to understand the mechanisms underlying binary choices^1,43^. One useful feature of these models is that they allow separation of different cognitive processes that drive choice. Here, we introduce a multi-attribute, time-dependent, DDM (mtDDM) which allows two distinct and often opposing attributes, taste and health, to be processed at different times and weighted differently in the decision process. We show that both the influence of an attribute on evidence accumulation and the delay before an attribute contributes to the evidence accumulation process differ significantly by attribute – and that between-attribute differences in these two parameters explain a large proportion of the variance in healthy choices. This indicates that models assuming the relative value signal reflects the total stimulus value – and not potentially independent attributions – may be unnecessarily limited in their explanatory power.

Poor dietary choices are often attributed to the combination of two factors: strong preferences for the tasty foods that are endemic to modern society, and limitations in how well self-control mechanisms can inhibit the strength of those preferences^44^. Our findings support the alternative explanation that tasty dietary choices reflect not only of relative strength of taste preferences but also their relative timing^13,14^. That is, an individual may eat a cookie not because the desire for a tasty snack overwhelms their limited willpower, but because information about future health consequences does not enter the decision process sufficiently early to influence choice. Hereafter, we explore the implications of our results both for models of the decision process and for understanding decision making in the face of competing goals.

Our findings have several implications. First, they generate the clear recommendation that slowing down the decision process may mitigate the effects of relative attribute latency or lower weighing of health, which could improve choices for some multi-attribute decisions. Further, this suggests a mechanistic explanation for previous work showing that the relative encoding of taste information in in value-related brain regions decreases when free response times are allowed and increases with shorter response times^45^, such as time pressure^46^, which alters parameters of the DDM^47^. Future interventions could either remove time pressure from dietary choices where they often occur, such as at a drive-through window, or extend the decision process by mandating a waiting time before choice.

Second, we find that a prime that explains the importance of healthy eating can decrease the weight placed on taste information during evidence accumulation, facilitating more healthy choices. Such primes can readily be incorporated into choice architectures, allowing future work to test variations of this prime and its application outside the lab – which may provide opportunities for improving choice^29^.

Third, we propose that the processes identified for simple multi-attribute dietary choices should exist for other decision domains in which values may be processed differently. For example, in financial choices, one must often make a trade-off between spending money now, and saving for the future^48–50^ – and, similar to what is seen for dietary choices, the future consequences of financial saving may not be as readily estimated as the immediate benefits of spending now. This may lead to a slowed estimation of the value of delayed financial rewards, and therefore more impulsive choices, regardless of an individual’s underlying preference for saving. Similarly, a multi-attribute DDM has been proposed for social decision making^15^, and adding a latency parameter could extend this work. For example, the speed with which rewards for the self and others are processed and incorporated into the decision process may increase the model’s explanatory power, as well as individual differences in prosociality. Applying the mtDDM to different choice contexts, and with different forms of nudges, could help expand our understanding of both the decision process and how to improve choice.

There are multiple limitations to the mtDDM that could be addressed in future studies. First, our model assumes that drift slopes begin at zero but transition discretely to some fixed weight following a latency period. However, many cognitive mechanisms could alter the drift slope over time. For example, attention has found to significantly influence the evidence accumulation^5^. Second, plausible alternative models exist, such as one with a stopping time for an attribute’s consideration. Although we show here that the mtDDM outperforms a stopping time DDM, there may be other choice problems for which it improves model predictions. A time-variable drift rate^51^ could address both alternatives by assessing how drift slopes vary over time; for example, such a model could be implemented for dietary choice by down-weighting the taste drift slope once health information is computed. In addition, the mtDDM presented here assumes that taste and health combine linearly to guide choice. However, non-linear utility functions are often more robust^52^. For example, in monetary decision making, a hyperbolic model is often used to combine immediate and future value information into a singular utility to guide choice ^48,53,54^. Future work could probe the precise functional form appropriate for evidence accumulation.

Moreover, previous work has found significant trial-to-trial variability in DDM parameters^36,55^. Examination of these fluctuations may help explain within-individual variability in dietary choice. For example, it is possible that when a healthy option is chosen, there is a reconsideration time that does not exist (or is different) for unhealthy choices and that could lead to a shorter health latency. Although we find that health’s latency is still longer, on average, in trials in which a healthy choice is not possible, and that latencies could be recovered even with such reconsideration times, examination of the various ways in which health information can enter the decision process would be a fruitful avenue for future research.

Another set of potential limitations are methodological: interdependencies can arise between parameters in multi-parameter models. For example, smaller drift slopes and larger boundary widths could produce similar choice and response time patterns. This is also a concern with the mtDDM, although our successful parameter recovery indicates parameters can be estimated with at least some accuracy. Additionally, the current work presents parameters estimated using only choices and response times. Although this is convenient (both are readily obtained via standard methods for both laboratory and naturalistic experiments) it is also a limitation. This work could be extended by including neural signals, which may provide more accurate estimates or refinements to the model itself. Previous work using neural data to inform multi-attribute choices and models^7,15,56^ are a promising direction.

Finally, our work suggests that different interventions may work better for some individuals than others. For example, individuals with very slow processing of health information might benefit most from extending their decision process by introducing a wait time before choice. For others who weigh health minimally or not at all in choice, extending decision time may not substantially improve choice; instead, interventions would need to first encourage consideration of health information (in any form) through a mechanism such as priming. By broadening interventions beyond appeals to self-control to include a more nuanced consideration of the timing and strength of different attributes, researchers and policy makers will be more likely to identify methods for eliciting healthy choices.

## Methods

All procedures and stimuli were approved by the Institutional Review Board at Duke University.

### Participants and sample size

Seventy-nine young adults from the Durham-Chapel Hill community (64% female; mean age 24.4 years) participated in this 90-minute study. Participants were screened for any dietary restrictions. Informed consent was obtained after the experiment was explained to participants.

The targeted sample size (40 individuals in each of two priming groups) was determined based on measurements in two independent datasets (results in preparation for publication) that included a binary choice task like our task described below. First, we calculated the effect of our differential priming conditions on the proportion of healthy choices across a large sample of subjects (N=133), which generated an approximate required sample size of between 40 and 45 participants in each prime group (via the sampsizepwr function in MATLAB and a *p*<0.05 threshold for effects by prime). We next examined the robustness of our priming effects in a second independent data set (N=40), in which the main effect of our primes fully replicated. Based on these prior results, we set 40 participants in each prime group as the target sample size in the current study.

One participant did not have sufficient variability in food ratings to generate 150 Conflict Trials; that participant is not included in analyses involving the proportion of healthy choices in Conflict Trials.

### Experimental procedure

Prior to the experiment, participants fasted for four hours, with compliance as measured by computerized self-report. Participants were compensated with $12 in cash and a snack food for consumption at the end of the experiment. All stimuli were presented with the Psychophysics Toolbox ^57^ for MATLAB. The experiment contained four phases, always presented in the below order. See Supplemental Methods for task instructions.

#### Phase 1: Rating Task

Participants began by rating 30 familiar snack foods on three five-point scales. They were asked their opinions of the tastiness, healthfulness, and wanting (“How much do you want to eat this food at the end of the experiment?”). Scale type, food presentation order, and left-right scale direction (good to bad, or bad to good) were randomized across participants. Stimuli were 600 x 600 pixel full-color images on a black background, presented alongside a one- to three-word item name (e.g. “Oreos”). Food images included a sample of the food outside of its packaging (e.g., a few chips outside the chips bag).

#### Phase 2: Goal Priming

Each participant was randomly assigned to one of two priming conditions. After the ratings task, participants read instructions for the following Food Choice Task (described below). A short instructional script (see Supplemental Methods) was imbedded in these instructions. This script emphasized the importance of either health information (“Health Prime”; N=40) or taste information (“Taste Prime”; N=39) in dietary choice using science-based reasoning. Data collection and analysis were not performed blind to the prime condition.

#### Phase 3: Choice Task

Next, participants made 300 self-paced choices between pairs of foods they had rated in Phase 1. On each trial, they saw two foods and indicated which they would like to eat more using a keyboard (Fig. 4a) and were told that one trial would be randomly selected, and that food would be served to them at the end of the experiment. Using the participants’ previous food ratings, half of the trials were constructed with one food that was tastier and less healthy than the other food (“Conflict Trials”). Note that one participant did not have enough variance in health and taste ratings to construct 150 Conflict Trials; for that participant, foods were paired randomly, and any reported statistic measuring the proportion of healthy choices made in Conflict Trials does not include this participant. One third of trials presented options using images, one third as their item names from the ratings task, and one third featured one option in words and the other as an image; as this study does not focus on differences in choice by image presentation, data from all three option representation trial types are pooled together to maximize the number of trials used for more precise parameter estimation. Presentation order was randomized across trials and participants, while ensuring that the same item did not appear within five trials.

Participants then completed a second version of the food choices task and personality questionnaires; those measures are outside of the scope of this paper and not reported here. The analyses reported here were not tested or performed on this second task, which was part of a larger series of tests of dietary nudges; this second task was always performed after the one used in these results, and participants were not aware that it would occur. For the results of this second task, see^58^.

#### Phase 4: Incentive Delivery

To ensure incentive compatibility, at the end of the experiment one trial was randomly selected, and the food chosen on that trial was given to the participant. Participants could leave immediately after eating one serving of the food or could wait thirty minutes in the experiment room (1 of the 79 participants chose to wait). This procedure encouraged participants to treat each trial as if it were the one that could count for their food compensation.

### Statistical Analysis

All statistical analyses were performed in MATLAB. All t-tests reported are two-sided. Data distributions were assumed to be normal, but this was not formally tested. All mixed effects regressions used mixed effects regressions with random slopes and intercepts using MATLAB fitglme. Between-subjects regressions were performed using MATLAB regstats. Pearson correlations were performed with MATLAB corr. Estimation of mtDDM parameters was performed using maximum likelihood estimation in MATLAB. Statistical thresholds were set to p<.05. All statistical tests that resulted in a p-value less than 0.001 are reported at that level, given the limits on the precision of our statistical analyses.

### The mtDDM

We simulated choices and response times for a multi-attribute, time-dependent DDM (mtDDM). In this model, a relative value signal (RVS) evolved in 10-ms time steps per convention. At each time step *t*, a weighted amount of the relative (left minus right) taste (T_L_–T_R_) and health (H_L_–H_R_) value difference was added the RVS. When the RVS reached the boundary for the right or left item, a choice was considered as being made for that food. The value signal evolved per equation (1). Parameter *τ* determines the drift latencies, set by t*_T_ and t*_H_:

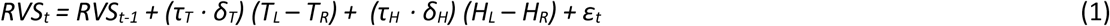

where,

*τ*_*T*_ = 1 if *t* ≥ *t*^∗^_*T*_, and *τ*_*T*_ = 0 otherwise;
*τ*_*H*_ = 1 if *t* ≥ *t*^∗^_*H*_, and *τ*_*H*_ = 0 otherwise.

In this model, ε represents i.i.d. Gaussian noise with a standard deviation fixed to σ = 0.1. The drift latency parameter t* represented the time before which each attribute’s relative value does not contribute to the RVS, and after which it contributed at a rate determined by its drift slope. For speed of estimation, this model assumes that the non-decision time proposed in standard DDMs (i.e., the time during the trial not allocated to evidence accumulation) is included in both taste and health drift latencies. One parameter commonly used in diffusion modeling is bias at choice outset, often resulting from over-trained motor response as it is introduced before options are identified or processed. As options in this task were randomly and equally presented on the left and right sides of the screen, participants were unable to develop a pre-set motor bias toward the healthier or tastier item on each trial. Further, choices in this mtDDM were fit using left vs. right choices, and not healthy vs. unhealthy choices. Therefore, bias was fixed to zero (i.e., in favor of neither the left nor right option).

### Per-participant DDM Parameter Estimation

We estimated five parameters of the mtDDM (taste and health drift slopes, taste and health drift latencies, and boundary width) for each participant in MATLAB. Using a multi-stage grid search, then optimized using nonlinear minimization. The best-fitting parameters for each subject were determined using maximum likelihood estimation. See Supplemental Information for more details on this procedure.

### Parameter Recovery

We performed a parameter recovery exercise of 100 simulated participants to ensure that simulated mtDDM parameters could be recovered using our estimation methods. See Supplemental Information for more details on this procedure and its results, and Figure S7 for correlation between true and recovered parameters.

### DDM Simulation to Illustrate Model Predictions

To generate the qualitative predictions for the influence of taste and health latencies on response times and choices displayed in Figure 2, a stimulus set was constructed using health and taste values like those in the experimental dataset’s conflict trials – specifically, all possible combinations of value differences ranging from −4 to 4 in which one option had a larger health, and smaller taste, than the other Taste’s drift slope and latency were fixed to 0.08 units/ms and 500 ms, respectively. Health drift slopes were varied between from .04 to .12 units/ms in .02 increments had health latencies that ranged from 10 ms to 1000ms, in 250 ms timesteps. Boundary size was fixed to 1 unit. For each of the 25 parameter combinations and 16 taste and health value difference pairs, 1,000 decision processes were simulated, and proportion of healthy choices and mean response times were recorded.

## Supplementary Information

### Supplemental Methods

#### Task instructions

##### Ratings Task

In the first task, you will rate foods on three attributes, one at a time. Ratings will be made using the keyboard. You’ll rate foods on their health, how tasty they are to you, and how much you would like to eat them at the end of the experiment. You have the option to rate an item as “neutral”, but please avoid that as much as possible. Rate each food based on how it looks on the screen. For example, when rating a picture of a plain piece of bread, tell us how healthful you think it is alone, not how healthful it would have been if it were covered in butter. For each food, you will be shown both an image and a text description. It is important to pay attention to both the image and its text description when rating the foods. This is because later in the experiment you may have to make decisions based on the text descriptions alone.

##### Choice Task

[Either Health or Taste Prime Text; see below]. On each trial of this task, you will choose between two different foods. Press the one (1) key to select the food on the left, and the zero (0) key to select the food on the right. Your choices will be displayed as either food images or their text descriptions. The descriptions will be the same text descriptions that you saw in the previous ratings task. At the end of the experiment, you will actually receive your food choice from one randomly-selected trial across the entire experiment (from either this task, or next task). You can leave either when you’ve eaten the food, or when one half hour has expired.

#### Prime text

Imbedded in the Choice Task instructions, participants received either a health or taste prime to encourage either health or taste goals in their subsequent binary choices.

##### Health Prime

The purpose of this study is to learn about how a food item’s healthfulness affects people’s choices about what they eat. We are interested in this question because leading scientists at top universities across the country have shown that eating a healthy diet is very important. They mention that one key benefit of eating healthy is the ability to maintain a healthy body weight, which can reduce the risk for many diseases. Previous research found that the top three killers in America are heart disease, cancer and stroke. Chronic diseases develop over time and are the cumulative effect of each eating decision we make in our lives. The health benefits of eating healthy are clear, but we would like to better understand how people incorporate health into each individual food choice.

##### Taste Prime

The purpose of this study is to learn how taste affects people’s choices about the foods they eat. We are interested in this question because food is a central part of human culture, and is thought to be a source of enjoyment, passion, and fulfillment for many. Leading scientists at top universities across the country have found that high-taste foods reliably increase activity in the brain’s reward centers. This increased activity is usually associated with a boost in dopamine levels in the brain, and dopamine is closely tied to our brain’s reward processing, as well as our subjective experiences of reward. We would like to understand how food choice is affected by the rewarding aspects of flavor and taste. The benefits of eating flavorful foods are clear, but we would like to better understand how people incorporate taste into each individual food choice.

#### mtDDM Model Estimation

In this model we estimated five free parameters: δ_T_, δ_H_ t*_T_, t*_H_, and b. Best-fitting mtDDM parameters were estimated separately for each participant using the following procedure. For each unique taste-health rating value pair experienced by the participant during the choice task, 5,000 simulations were run for each mtDDM free parameter combination to compute the likelihood function over observed left or right item choices and their response times. All trials with response times less than 8000 ms and greater than 300 ms were used, and response bins were created using MATLAB’s histcounts. The best-fitting parameter set was found by selecting the set that resulted in simulated choices and response times that minimized the negative log-likelihood of the experienced trials.

Parameter estimation was run in four stages. Stage one was used to find the maximum and minimum search values for each free parameter. This was done by running a coarse grid search for each participant and increasing maximum, and lowering minimum, search values until no participants hit the maximum or minimum value for any parameter. Based on this procedure, the maximum and minimum parameter values for each attribute (taste or health) to search over for stage two were set to: drift slope, [max(min(δ) - 0.1, 0), max(δ) + 0.1], drift latency, [max(min(t*) − 500, 10), max(t*) + 500]. Boundary width, which did not differ by attribute, was set to [max(min(b) − 0.2, .1), max(b)+0.2]. Noise added to each time step was set to be normally distributed around zero with a standard deviation of 0.1.

Stage two served to find a finer estimate of each participant’s free parameters. The final grid, after expanding the maximum and minimum values as described above, consisted of every possible combination of the following parameters: Drift slope for taste, δ_T_, [.02, .16] in .01 increments. drift slope for health, δ_H_, [0, .16] in .01 increments, boundaries, b, [.4, 2] in .1 increments. Drift latency for taste, t*_T_, [10,2240], and for health, t*_H_, [10, 2230], both in 50 ms increments. To speed estimation, the subject’s maximum possible drift latency was re-set to the average response time for that participant if it was smaller, as a latency longer than a mean response time is not physiologically meaningful.

The purpose of the third stage was to find the best-fitting set of parameters at a much finer resolution. To do this, the best-fitting stage two parameters for each participant were used to develop a very fine search grid. The grid for each parameter was set using the following procedure. For drift slope, the upper and lower grid values were +/− .01 the stage two estimates in .001 increments, excluding δ<0; for boundaries, the upper and lower values were +/− .01 of the stage two estimate, in .01 increments; for both t*_T_ and t*_H_, the upper and lower grid values were +/− 500 ms, in 10 ms increments, excluding t* < 10 ms.

The fourth stage used a nonlinear minimization search using MATLAB’s fmincon to search within stage three’s fine grid to arrive at the final parameter combination for each participant. For each participant, the top five best-fitting parameter combinations from stage three were selected. For each of these five-best combinations, the minimization was run 1,000 times, resulting in a total of 5,000 parameter estimations. The starting point for each parameter on each of these 5,000 minimizations was drawn randomly from a uniform distribution whose median was that parameter’s stage three best-fitting value, and whose maximum and minimum values were that best-fitting value plus and minus stage three’s step size. These maximum and minimum values were also the maximum and minimum constraints on each parameter’s search. This effectively allowed for a random search within the best-fitting grid from stage three.

For all stages, there was only one best-fitting parameter combination for each participant (i.e., only one smallest negative log likelihood). The best-fitting parameters from the fourth stage were used for the analyses reported in the paper.

#### Single-Latency Model (mDDM)

We next tested the ability of the mtDDM to explain choices and response times better than a model without taste and health latency parameters. To do this, we estimated a DDM model identical to the mtDDM, but that assumed that both taste and health enter the decision process at the same time (See Supplemental Materials for details), which we term “value latency.” We fit separate taste and health drift slopes. The model was as follows:

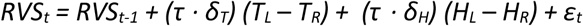

where,

*τ* = 1 if *t* ≥ *t*^∗^, and *τ* = 0 otherwise.

#### Attribute Stopping Time Model (stDDM)

We tested a model in which we replaced attribute latency parameters with attribute stopping time parameters. In this model, both attributes begin contributing to the decision process at the same time, but the time at which they stop contributing was allowed to vary. This was estimated using the same procedure described above. The model was as follows:

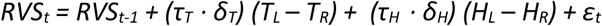

where,

*τ*_*T*_ = 1 if *t* ≤ *t*^∗^_*T*_, and *τ*_*T*_ = 0 otherwise;
*τ*_*H*_ = 1 if *t* ≤ *t*^∗^_*H*_, and *τ*_*H*_ = 0 otherwise.

#### Latency Only Model (latDDM)

We tested a model in which both taste and health receive the same drift slope δ, which is a temperature parameter. Taste and health latencies were allowed to vary. This was estimated using the same procedure described above. The model was as follows:

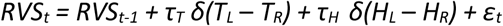

where,

*τ*_*T*_ = 1 if *t* ≤ *t*^∗^_*T*_, and *τ*_*T*_ = 0 otherwise;
*τ*_*H*_ = 1 if *t* ≤ *t*^∗^_*H*_, and *τ*_*H*_ = 0 otherwise.

#### Simple DDM (sDDM)

We tested a simple DDM with only three parameters, in which taste and health receive the sample slope drift slope δ and taste and health latencies held to be equal. This was estimated using the same procedure described above. The model was as follows:

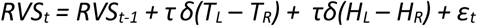

where,

*τ* = 1 if *t* ≤ *t*^∗^, and *τ* = 0 otherwise;

#### Parameter recovery procedure

Choices and response times were simulated with the same mtDDM process used for parameter estimation. mtDDM parameter combinations were randomly selected from the range of values in our participant dataset. For each of 100 agents, a dataset of 5,000 trials were simulated using all possible taste and health value difference combinations (from −4 to 4) in the experimental dataset. On half of those trials, to approximate the experimental dataset’s composition of trials, health and taste were in conflict (that is, one option was tastier, and less healthy, than the other) Next, the same parameter estimation procedure applied to the participant data was used to estimate the five free mtDDM parameters (see “mtDDM Model Estimation” above).

#### Cross-validation DDM fitting

To compare DDM parameters to behavioral measures such as choices and logistic decision weights while avoiding “double dipping” (comparing parameters to choices used to fit the parameters), a separate set of DDM parameters was fit for each participant using a cross-validation method. First, one half of a participant’s 300 choices were randomly selected for parameter estimation. This was done such that half of a participant’s non-conflict trials, and half of their conflict trials, were used. This resulted in a dataset for each participant of 150 total trials, 75 conflict and non-conflict trials each – except for the participant with no conflict trials, in trials were partitioned randomly. DDM parameters were then fit using the procedure detailed above (“mtDDM Model Estimation”). For all analyses performed with the cross-validated parameters, comparisons were made between the fitted parameters from one half of trials, and choices on the other half of trials (that is, those trials not used to estimate the DDM parameters).

### Supplemental Results

#### mtDDM Predictions Using Simulations

We first assessed how relative value of the healthy vs. unhealthy option (in this simulation, value was the unweighted sum of taste and health) was related to healthy choice, and how taste and health latencies influenced this relationship. This was done with a series of simulations, in which mtDDM parameters were used to predict the proportion of healthy choices made. In this analysis, health and taste drift slopes and boundary size were held to be equal. Taste latencies were fixed, while health latencies were allowed to vary. This allowed us to see, qualitatively, how changes in the relative latency of health would influence the proportion of healthy choices made, holding all else equal (see Supplementary Methods above).

In Figures S2a and b, the black line refers to simulations in which taste and health latencies were equal. We found that as health latencies became faster (Fig. S2a, from dark to light blue), the proportion of healthy choices increased as a function of value, as evidenced by a leftward shift in the psychometric choice curve. Conversely, the psychometric curve shifts to the right as health latencies became slower (in red), indicated an increase in unhealthy choices and taste latencies become relatively faster. The simulations also predict a pattern in response times that would be expected from value-based choices: specifically, that response times will be longer when both options are similar in value (Fig. S2b).

Figure S2c displays the simulation in a different way to highlight specifically the proportion of healthy choices predicted (y-axis) relative to health’s latency (x-axis), which was varied in these simulations. Importantly, this also demonstrates how this relationship is also influenced by the value difference between the options. As mentioned above, taste latency was always fixed to 200 ms (vertical grey bar). As health latencies became faster than taste’s (to the left of the grey bar), the proportion healthy choices increased - even when the value of the healthy item was much worse than the tasty item (blue lines). Healthy choices declined as health’s latency became increasingly slower than taste’s latency or slower than (to the right of the grey bar). Of note, latency had an influence on choice even when one option was strongly preferred to the other (value differences of - 4 in blue, or 4 in green), and that this influence is strongest when both options are equally liked (light blue, value difference of 0). This further illustrates that earlier health latencies predict more healthy choices, even in cases in which the tempting tasty item was higher in value.

#### Response Times by Choice and Trial Type

Response times (RTs) were faster for conflict trials (in which there was a choice between a healthier, less tasty food and a tastier, less healthy food) than in nonconflict trials (Fig S3a; M = 1558 ms, 1636 ms; paired t-test of log(RTs) d = −0.17, t(77) = −3.34, p = 0.001, 95% CI = [−0.08 −0.02]). Among conflict trials, trials in which the healthier choice was made took longer (Fig S3b; M = 1917 ms, 1493 ms; paired t-test of log(RTs) d = 0.61, t(76) = 7.33, p < 0.001, 95% CI = [0.15 0.27]). There was not a statistically significant relationship between the number of healthy choices made and RTs when making healthy choices (Pearson correlation ρ = −0.02, p = 0.86). However, participants who made more healthy choices took longer on unhealthy choice trials (Pearson correlation ρ = 0.26, p = 0.02). This is expected from any DDM with separate weights on taste and health and is the result of compounding of the weight placed on health versus taste during the decision process, a prediction we pursue below.

##### DDM Predictions for RT Differences Between Choice Types

Both the mDDM and mtDDM would predict the observed phenomena of slower healthy choices and faster unhealthy choices, given that individuals (on average) place a larger weight on taste than health in the decision process. To confirm this analytically, we performed a series of simulations to 1) assess whether a multi-attribute DDM can predict the observed RT differences between healthy and unhealthy choices, and 2) whether adding an attribute-wise latency parameter improves the prediction of these RT differences (above a multi-attribute DDM with one latency parameter). To answer these questions, we simulated RTs on each experienced conflict trial (in which one food was healthier and less tasty than the other). For each trial, 1,000 decision processes were simulated twice: Once using participants’ best-fitting mtDDM parameters, and once using participants’ best-fitting mDDM parameters (that is, a model identical to the mtDDM but fit with a single latency). We then compared the simulated RTs between healthy and unhealthy choice trials. For cases in which a simulation predicted no healthy or unhealthy choices, that trial type is omitted from the analysis.

Both the mDDM and mtDDM predicted longer RTs for healthy than unhealthy choice trials (mDDM M = 2101 ms, 1836 ms, paired t-test of log(RTs) d = 0.64, t(30) = 5.33, p < 0.001, 95% CI = [0.05 0.12]; mtDDM M = 2117 ms, 1804 ms, paired t-test of log(RTs) d = 0.71, t(31) = 6.00, p < 0.001, 95% CI = [0.10 0.20]). This is expected, as trials in which the healthier option is chosen are more likely to have smaller summed weighted attribute values due to the small weight on health (i.e., small drift slopes), which will result in relative value signals that take longer to cross a boundary. As the healthier option approaches and overtakes the unhealthy option in summed weighted value, two things happen: first, the relative value signal is likely to take longer to cross a boundary due to more similar value differences and second, the choice is more likely to be in favor of the healthy option. For example, consider a participant with drift slopes of taste=0.02 and health=0.01. For choice in which the relative taste and health of the (healthy - unhealthy option) is [taste = −4, health = 4], this results in a total weighted summed value of 0.02*−4 + 0.01*4 = −0.04, which is in favor of the unhealthy option. For a value difference pair, in which the healthier option has improved in its relative tastiness, of [taste = −1, health = 4], the total value is 0.02*−1 + 0.01*4 = 0.02, which is a smaller relative value difference but now in favor of the *healthy* option. The latter case will result in slower choices in favor of the healthy option and is an illustration of how healthy choices are often slower, as their summed weighted value is often much less due to the smaller weight on the health attribute. Next, we assess whether multi-attribute latencies improve explanation of this difference in RT between choices.

We investigated where specifically the mtDDM outperformed the mDDM by investigating RT and choice predictions by RT quantile. We note, however, that these comparisons do not penalize the mtDDM for its additional parameter, so therefore should be treated as exploratory. The mtDDM predicted larger differences in RTs between healthy and unhealthy choices than mDDM (healthy – unhealthy RTs, 170 ms vs. 286 ms; paired t-test of log(RT) differences d = −0.55, t(24) = −3.24, p = 0.003, 95% CI = [−0.09 −0.02]). This indicates that both models predict an RT difference, and that the mtDDM predicts an even larger difference between choice types. Next, we assess how well both models’ predictions match with the observed RT data.

Observed RTs were correlated with those predicted by both the mDDM and mtDDM but were more closely correlated for the simulated RTs of the mtDDM than the mDDM. The simulated mDDM and mtDDM RTs and observed RTs were correlated for both healthy (mDDM, ρ = 0.66, p < .001; mtDDM, ρ = 0.72, p < 0.001) and unhealthy (mDDM, ρ = 0.88, p < 0.001; mtDDM, ρ = 0.91, p < .001) choices.

Finally, we measured how well the DDM models predicted *differences* in RTs between healthy and unhealthy choices. For this, too, the mDDM does well, and the mtDDM does best. The difference in RTs between healthy and unhealthy choices are correlated between simulated and real RTs for both the mDDM and mtDDM (mDDM, ρ = 0.39, p = 0.005, mtDDM ρ = 0.62, p < 0.001). This correlation is larger for mtDDM than mDDM predictions (Williams-Hotelling test t(76)=3.74, p<0.001).

We further investigated RT predictions for different decision durations. For each participant, we binned experienced trials by log(RT) quantile and then used both their mDDM and mtDDM fitted parameters to predict RTs for each experienced trial. We found that the variable latency allowed by the mtDDM improves its predictions over the mDDM for the first three quantiles; interestingly, for the longest RT quantile both models have worse performance, with the mDDM providing better prediction.

###### Percent RT quantiles predicted correctly, mtDDM vs. mDDM

Q1 Mean difference = 0.058, d = 0.01, t(78) = 0.10, p = 0.920, 95% CI = [−1.089 1.205]
Q2 Mean difference = 2.950, d = 0.24, t(78) = 2.15, p = 0.035, 95% CI = [0.213 5.688]
Q3 Mean difference = 3.291, d = 0.15, t(78) = 1.33, p = 0.094, 95% CI = [−1.737 8.319]
Q4 Mean difference = −5.700, d = −0.26, t(78) = −2.33, p = 0.022, 95% CI = [−10.561 −0.836]

Similarly, the mtDDM does best relative to the mDDM in the 3^rd^ quantile, in predicting the correct choice. Comparing this performance by a median split of absolute differences in latency, we find that difference in mtDDM and mDDM performance is significantly better for the mtDDM for those with larger median latency differences in Quantile 3 (Means = 2.96, 0.94, d = 0.43, t(77) = 2.04, p = 0.044, 95% CI = [0.052 3.985]).

###### Percent Choices predicted correctly, mtDDM vs. mDDM, by RT quantile

Q1 Mean difference = 0.081, d = 0.02, t(78) = 0.19, p = 0.847, 95% CI = [−0.751 0.914]
Q2 Mean difference = 0.772, d = 0.24, t(78) = 2.17, p = 0.033, 95% CI = [0.063 1.480]
Q3 Mean difference = 1.941, d = 0.43, t(78) = 3.85, p < 0.001, 95% CI = [0.938 2.944]
Q4 Mean difference = 0.945, d = 0.20, t(78) = 1.77, p = 0.081, 95% CI = [−0.118 2.008]

#### mtDDM Simulation and Recovery

To ensure that no parameter of the mtDDM could be estimated to be falsely larger or smaller than its true value due to an inability to separate and estimate each parameter individually, or a bias causing artificial correlation between parameters, we assess how well each parameter of the mtDDM could be recovered from simulated data (see “Parameter recovery procedure” above for methods).

To assess the ability of this method to accurately recover the true parameters, we first assessed whether the recovered and true parameters were correlated, even if the precise parameter values were not recovered. There was a statistically significant correlation between true and recovered values for all parameters (scatter plots, Fig. S5; Pearson correlations: Drift Slope 1, ρ = 0.75, p < .001; Drift Slope 2, ρ = 0.71, p < .001; Drift Latency 1, ρ = 0.82, p < .001; Drift Latency 2, ρ = 0.87, p < .001; Boundary Width, ρ = 0.86, p < .001). Despite this, slopes and boundaries were recovered to be larger than their true values, and latencies were recovered to be shorter (histograms, Fig. S5; Recovered – True value mean differences, Drift Slope 1 = 0.003, d = 0.61, t(99) = 6.07, p < 0.001, 95% CI = [0.002 0.003]; Drift Slope 2 = 0.003, d = 0.59, t(99) = 5.87, p < 0.001, 95% CI = [0.002 0.004]; Latency 1 = −140 ms, d = −0.31, t(99) = −3.08, p = 0.003, 95% CI = [−230 −50]; Latency 2 = −129, d = −0.29, t(99) = −2.94, p = 0.004, 95% CI = [−215 418]; Boundary Width, = 0.15, d = 0.91, t(99) = 9.05, p < 0.002, 95% CI = [0.12 .18]).

As much of our hypothesis revolved around the *relative* attribute slope and latency (Taste – Health), we also assessed the relationship between the true and recovered relative drift parameters. Importantly, there was a statistically significant correlation between the true and recovered relative drift slope (ρ = 0.85, p < .001), and between the true and recovered relative drift latency (ρ = 0.93, p < .001). The difference between the true relative (Taste – Health) and recovered relative value was not statistically significantly different from zero for slope or latency (relative slope mean difference = −0.0002, d = 0.03, t(99) = 0.25, p = 0.80, 95% CI = [−0.0011 0.0014]; relative latency mean difference = 11.1 ms; d = 0.02, t(99) = 0.19, p = 0.85, 95% CI = [−103 125]).

We also use the mtDDM parameter recover to test for a major concern for parameter estimation. Specifically, it is possible that a large drift slope for one attribute could lead to recovering that attribute’s latency as faster than it truly was. To see if this happened in the simulated dataset, we investigated cases in which an attribute had a larger drift slope relative to the other attribute, but a slower attribute latency. We calculated the proportion of simulated agents in which this attribute’s latency was falsely recovered to be faster instead of slower. Since, in this simulation, taste and health are arbitrary attributes, we collapse across both attributes for this test. Because parameter combinations were made randomly, this happened half (50) of our 100 agents, compared to our participant data in which this happened for an attribute only 14% of the time for either attribute. For the simulated agents, a latency was incorrectly recovered as faster for 4 of the 52 agents, for a 92% accurate classification rate overall. Altogether, this exercise indicates that although it is possible for a larger drift slope to result in a falsely faster drift latency, it is unlikely that this happens systematically due to mis-estimation of mtDDM parameters.

#### Attribute Stopping Time DDM (stDDM)

We compared the mtDDM to a model in which both taste and health enter the decision process simultaneously, but stop being considered at different times. The same procedure was used to fit this model as the mtDDM. Once the stopping time for an attribute was reached, its influence on the value signal dropped to zero (that is, it no longer contributed to changes in the relative value signal over time).

As expected, drift slopes for the stDDM were larger for taste than for health in this model (M = 0.045, 0.015; d = 1.97, t(78) = 9.78, p < 0.001, 95% CI = [0.02 0.04]) and each attribute’s drift slope was correlated with its slope estimated using the mtDDM (taste, ρ = 0.86, p < 0.001; health, ρ = 0.45, p < 0.001). We found no statistically significantly difference in health slope between this DDM and mtDDM (d = −0.14, t(78) = −1.19, p = 0.24, 95% CI = [−0.01 0.00]), whereas taste slopes were statistically significantly larger in the mtDDM (d = −0.67, t(78) = −9.35, p < 0.001, 95% CI = [−0.02 −0.01]). Fitted boundary widths were statistically significantly larger in the stDDM than mtDDM (M = 1.50, 1.41; d = 0.40, t(78) = 7.82, p < 0.001, 95% CI = [0.06 0.10]). The best-fitting taste stopping time was significantly earlier for health than for taste (M = 4145 ms, 2551 ms; d = 0.85, t(78) = 5.26, p < 0.001, 95% CI = [991 2198]). Interestingly, the taste stopping times exceeded mean RTs for 96% of participants (vs. 58% for health stopping times), suggesting for most trials and participants, taste remained influential on the decision process until the decision boundary was reached.

Next, we assessed whether the addition of attribute-wise stopping time parameters improved the mDDM’s and mtDDM’s ability to explain choices and RTs. The mDDM resulted in statistically significantly smaller BIC values than the stDDM, indicating that the mDDM fit the observed data better (Table S2; paired t-test of BIC values, M = 1143, 1645; d = −4.65, t(78) = −58.50, p < 0.001, 95% CI = [−518 −484]). The mtDDM also had statistically significantly lower BIC values than the stDDM (paired t-test of BIC values, M = 1111, 1645; d = −4.99, t(78) = −63.30, p < 0.001, 95% CI = [−550 −517]).

Finally, we assessed whether stopping times can help explain any variance in healthy choices above drift slopes for taste and health. In a regression model predicting the percentage of healthy choices made by a participant, stDDM all parameters together explain a large proportion of choice variance (Table S3; R^2^ = 0.76, F(72) = 46.77, p < 0.001). As in the mtDDM, larger taste drift slopes were statistically significantly negatively related to the number of healthy choices (slope = −7.28, 95% CI = [−8.75 −5.80], p < 0.001) and health drift slopes were statistically significantly positively related to the number of healthy choices made (slope = 4.58, 95% CI = [2.91 6.26], p < 0.001). Later health stopping times were statistically significantly related to more healthy choices (slope = 3×10^−5^, 95% CI =[1×10^−5^ 4 ×10^−5^], p < 0.001). We found no statistically significant relationship between taste’s stopping time and healthy choices (slope = −7×10^−6^, 95% CI = [2×10^−5^ 1×10^−5^], p = 0.41), perhaps because stopping times exceeded the mean RT for 96% (all but three) participants and therefore had little influence on most choices.

#### Latency DDM (latDDM)

We compared the mtDDM to a model in which both taste and health enter the decision process at different times, but their drift slopes are equal. The same procedure was used to fit this model as the mtDDM. Fitted latencies for taste were earlier than for health (M = 231.52, 1238.86; d = −1.97, t(78) = −10.46, p < 0.001, 95% CI = [−1199.01 −815.68]). Taste and health latencies estimated from the latDDM were correlated with those estimated in the mtDDM (taste, ρ = 0.71, p < .001; health, ρ = 0.55, p < .001). Taste latencies were estimated to be earlier than those estimated by the mtDDM (M = 231.52, 407.35; d = −0.58, t(78) = −6.55, p < 0.001, 95% CI = [−229.29 −122.37]), while health latencies were estimated to be later than those estimated by the mtDDM (M = 1238.86, 880.44; d = 0.63, t(78) = 5.74, p < 0.001, 95% CI = [234.02 482.81]).

Next, we calculated whether choices and RTs were explained better for the latDDM, mDDM, stDDM, or mtDDM using pairwise comparisons of Bayesian Information Criterion (BIC) values, which penalize additional drift parameters. The mtDDM resulted in statistically significantly smaller BIC values than the latDDM, indicating that the mtDDM fit the observed data better (Table S2; paired t-test of BIC values, d = −0.57, t(78) = −8.06, p < 0.001, 95% CI = [−69 −41]). This was also true for the mDDM (d = −0.23, t(78) = −3.00, p = 0.004, 95% CI = [−38 −8]). The latDDM fit the data better, however, than the stDDM (d = 4.51, t(78) = 49.97, p<0.001, 95% CI = [459 497]) and the sDDM (see sDDM details below; M = 1168.99, 1245.58; d = −0.83, t(78) = −8.40, p < 0.001, 95% CI = [−94.74 −58.44]).

#### Simple DDM (sDDM)

We compared the mtDDM to a model in both drift slopes and drift latencies for taste and health are fixed to be equal, which is functionally equivalent to a simple single-attribute DDM. Below we report comparisons between the fit of the sDDM to the other DDM models, which demonstrate that models with multi-attribute latencies perform better than the sDDM, but that models without them perform worse than the sDDM. The mtDDM fits the observed data better as measured by smaller BIC values (M = 1111.21, 1245.58; d = −1.44, t(78) = −12.97, p < 0.001, 95% CI = [−155.00 −113.75]), as does the latDDM (M = 1168.99, 1245.58; d = −0.83, t(78) = −8.40, p < 0.001, 95% CI = [−94.74 −58.44]). However, the sDDM has lower BIC values than the stDDM (M = 1644.45, 1245.58; d = 3.88, t(78) = 32.48, p < 0.001, 95% CI = [374.42 423.32]) and the mDDM (M = 1432.47, 1245.58; d = 1.53, t(78) = 10.81, p < 0.001, 95% CI = [152.47 221.31]).

#### Consideration of choice-specific non-decision times (csNDTs)

An alternative decision process, whereby participants reconsider their choice after a decision boundary is reached and that this time differs between healthy and unhealthy choices, could possibly bias latency estimation. Specifically, longer healthy choice non-decision times could lead to health latencies falsely estimated as shorter (and vice versa in unhealthy choice trials), and therefore a false difference between health and taste latencies. We address this possibility with three analyses. First, we note that the imbalance in healthy and unhealthy choice trials in our dataset makes it ill-suited to directly estimate choice-specific non-decision times. Participants made on average 33 healthy choices (range, 0 to 140) and 115 unhealthy choices (range, 0 to 150). This would lead to noisier healthy choice nondecision time (NDT) estimates than unhealthy choice NDT estimates, and also would mean that participants with many healthy choices would have more accurate estimates for their healthy-choice than unhealthy-choice NDT, and vice versa for participants making many unhealthy choices. Future work with an adaptive design to ensure an equal number of healthy and unhealthy choice trials is a promising avenue for future work.

##### Non-conflict trial mtDDM estimation

If attribute latencies are a relatively stable feature of the individual’s decision process, participants should have similar attribute processing times if estimated from non-conflict trials only (that is, trials in which one option is not healthier and less tasty than the other; half of all trials). These non-conflict trial latencies should predict healthy choice in the conflict trials. Because nonconflict trials do not have the option of a healthy choice, it would be impossible to have the csNDTs (as there are no healthy and unhealthy choices). To address this, we estimated mtDDM parameters for each participant using their non-conflict trials only. For the one participant who had only non-conflict trials, 150 trials were randomly selected for this estimation. We found that taste drift slopes were larger than health drift slopes (M = 0.05, 0.02 units/ms; d = 1.43, t(78) = 8.04, p < 0.001, 95% CI = [0.02 0.04]), and their difference was related to relative (taste – health) decision weights in conflict trials (Pearson ρ = 0.77, p < 0.001). Crucially, taste latencies were faster than health latencies (M = 342, 609 ms; d = −0.94, t(78) = −6.25, p = 0.001, 95% CI = [−351 −182]), and their difference also was correlated with relative decision weights (Pearson ρ = −0.25, p = 0.03). This indicates that, even when there can be no csNDTs specific to healthy and unhealthy choices, latency findings replicate those found with the entire trial dataset, and are associated with the healthy choices a participant makes when confronted by a conflict trial. However, we note that there were very few trials used for estimation, so these results should be treated with caution.

##### mtDDM parameter recovery using csNDT data generation process

We next used simulations and parameter recovery to test whether longer healthy choice NDTs would falsely recover as longer taste (vs. health) latencies (and vice versa for unhealthy choice NDTS). We did this by simulating choices and RTs as described in our mtDDM recovery with two changes. First, all trials were “conflict trials” so that all choices would be either healthy or unhealthy choices. Second, NDTs varied by healthy or unhealthy choice instead of by attribute. Then, mtDDM parameters were estimated from this data using the estimation procedure described above. This allows us to test whether a data-generating process with choice-specific and not attribute-specific latencies could result in differences in attribute-specific latencies (even though they were not a part of the decision process). Waiting times ranged from 100 to 500 ms in 10 ms timesteps which was chosen *a priori* informed by the range of non-decision times in our dataset (estimated from the single latency mDDM model which had a mean of 280 ms and within one standard deviation ranged from 101 ms to 468 ms).

First, we note that true and recovered drift slopes and boundaries were correlated (Taste slope, Pearson ρ = 0.82, p < 0.001; Health Slope Pearson ρ = 0.77, p < 0.001; boundary Pearson ρ = 0.56, p < 0.001). However, true choice-specific NDTs were not correlated with estimated attribute latencies (Fig. S9A; Taste latency vs. Healthy Choice NDT, Pearson ρ = 0.09, p = 0.38; Health latency vs. Healthy Choice NDT, Pearson ρ = 0.01, p = 0.90; Taste latency vs. Unhealthy Choice NDT, Pearson ρ = 0.00, p = 0.98; Health latency vs. Unhealthy Choice NDT, Pearson ρ = 0.18, p = 0.08). This indicates that there is no credible evidence to support the hypothesis that longer csNDTS would lead to different attribute latencies. We do highlight, however, that there was a non-significant trend toward a correlation between longer health latencies and longer unhealthy choice NDTs (p = 0.08) in the direction we expected if a bias did exist.

##### mtDDM parameter recovery using mtDDM and csNDT data generation process

For completeness, we performed a further simulation. This estimation was identical to the above, with one addition: taste and health latences were added to the decision process. This resulted in a DDM model with seven parameters: taste and health drift slopes, healthy and unhelathy choice NDTs, taste and health latencies, and boundary width. We then performed the same parameter recovery exercise to test whether attribute latencies be recoverd (that is, separately estimated) if both attribute-wise latencies and choice-specific NDTs exist in the decision proces. We find that, even when csNDTs are included, recovered taste and health latencies are correlated with their true parameters (Fig. S9B; Taste latency, Pearson ρ = 0.62, p < 0.001; Health latency, Pearson ρ = 0.63, p < 0.001, taste – health latency, Pearson ρ = 0.61, p < 0.001). However, we note that this is a noisier recovery than one without csNDTS (mtDDM recovery reported above), and that both taste and health latencies are recovered to be faster than their true values (one-sample t-test of difference between recovered and true parameters vs. 0; taste latency, M = 180 ms; d = 0.38, t(99) = 3.81, p 0.001, 95% CI = [87 274]; health latency, M = 223; d = 0.46, t(99) = 4.61, p < 0.001, 95% CI = [1267 319]). However, this indicates that taste and health latencies can be recovered fairly accurately even if csNDTs exist in the data-generation process.

Across three analyses, we find no credible evidence to support a concern that choice-specific non-decision times result in a large bias in attribute latencies and are therefore not likely to be a cause of significant bias in our manuscript’s results.

## Supplementary Figures

**Supplementary Figure 1.**
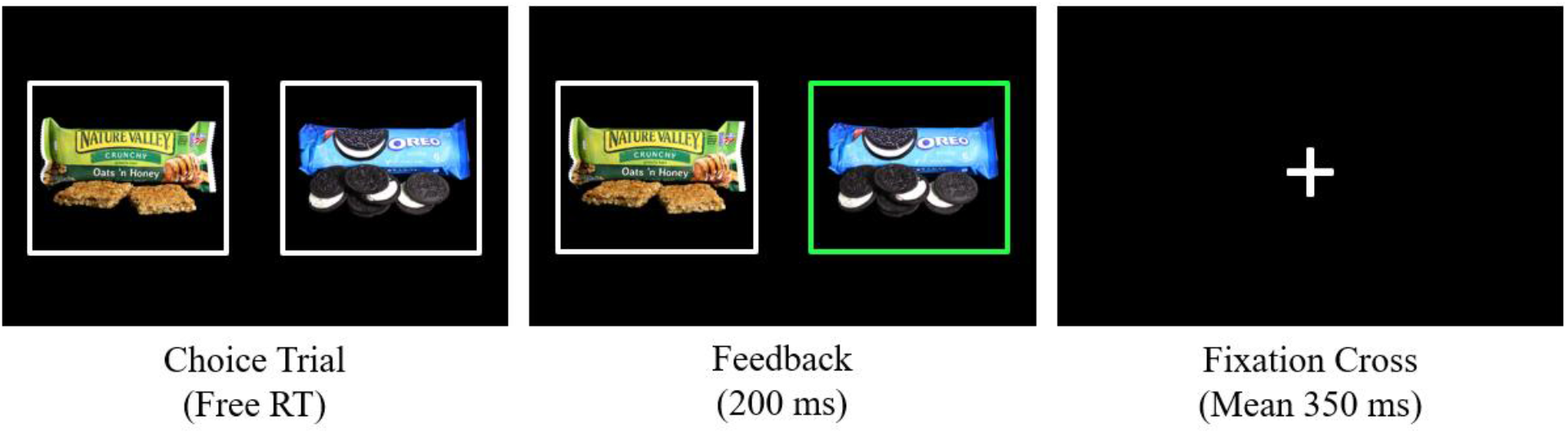
Flow of a choice trial. Participants made 300 binary choices with free response time (RT). Foods were displayed on the left and right sides of the screen. After keyboard response, the chosen food was highlighted in green for 200 ms to reflect participant response. Between trials, a fixation cross was displayed in the center of the screen for between 200 and 500 ms (mean 350ms; i.i.d. distributed).

**Supplementary Figure 2.**
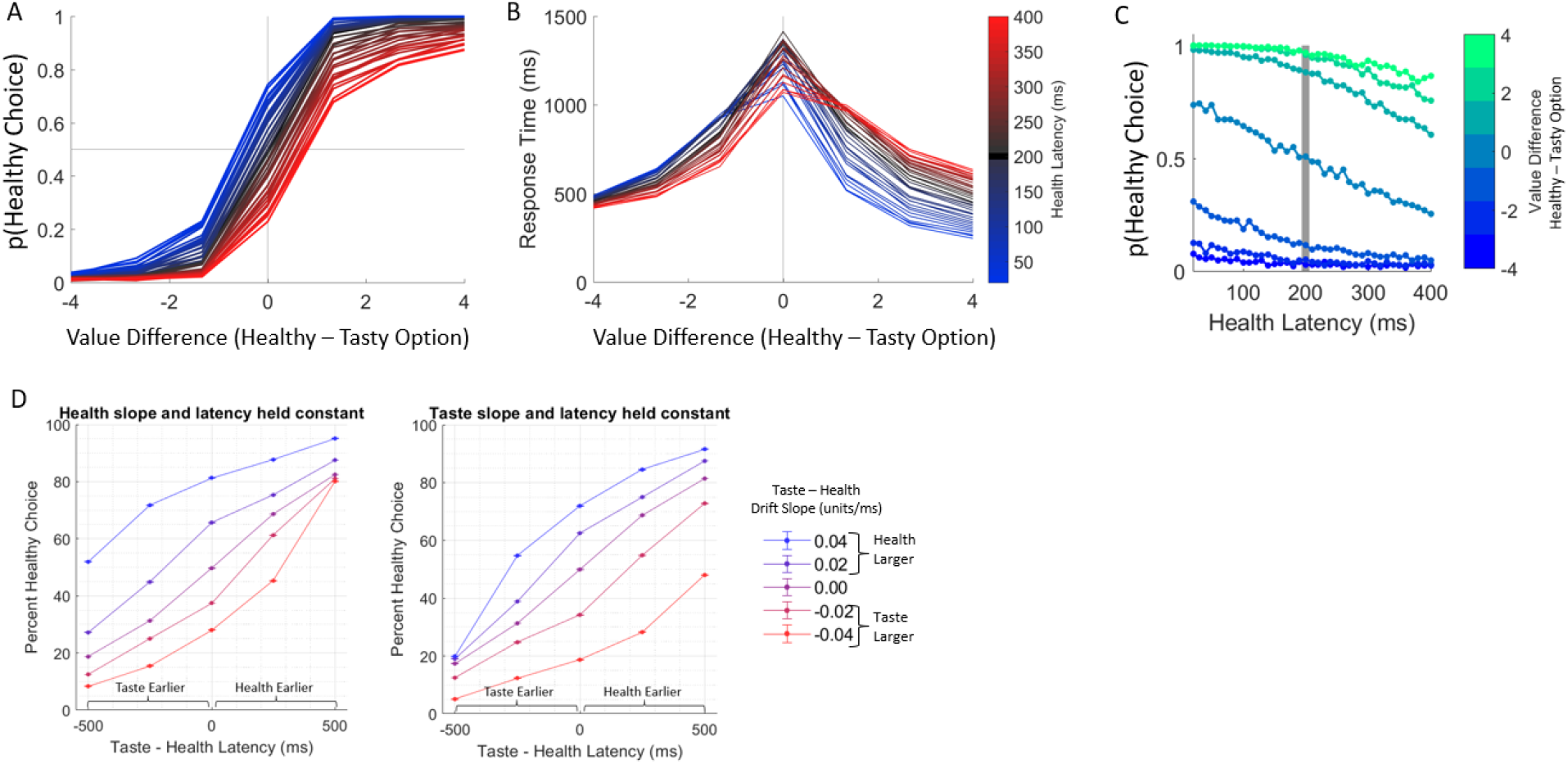
Simulation results. (**a**) The proportion of healthy choices for each value difference (Healthy– Tasty Food Value) are plotted for different relative latencies. Darker lines represent more similar latencies. Blue lines represent earlier health than taste latencies, and red lines represent earlier taste than health latencies. (**b**) Mean response times for each value difference (Healthy– Tasty Food Value) are plotted for different relative latencies; line colors are similar to those in the previous panel. (**c**) The proportion of healthy choices (y-axis) predicted as a function of health latency (y-axis) and option value difference (green to blue). When taste and health’s latencies are equal (at 200 ms, indicated by the grey bar), both attributes entered the decision process at the same time. Value differences, indicated here by colors green to blue, represent the relative (Healthy– Tasty) value (taste + health) of the options, with green representing a higher valued healthy option and blue representing a higher valued tasty option. As the healthy option increased in value (green), it was selected more often. There was a further increase in healthy choices moving from the grey bar to the left, indicating faster health latencies relative to taste latencies. Conversely, as health latencies became slower, plotted here to the right of the grey bar, fewer healthy choices were made. (d) The simulated percent of healthy choices predicted by the mtDDM given changes in taste slopes and latencies, while holding health (left panel) or taste (right panel) latencies fixed to 500 ms and health (left panel) or taste (right panel) slope fixed to 0.08 units/ms.

**Supplementary Figure 3.**
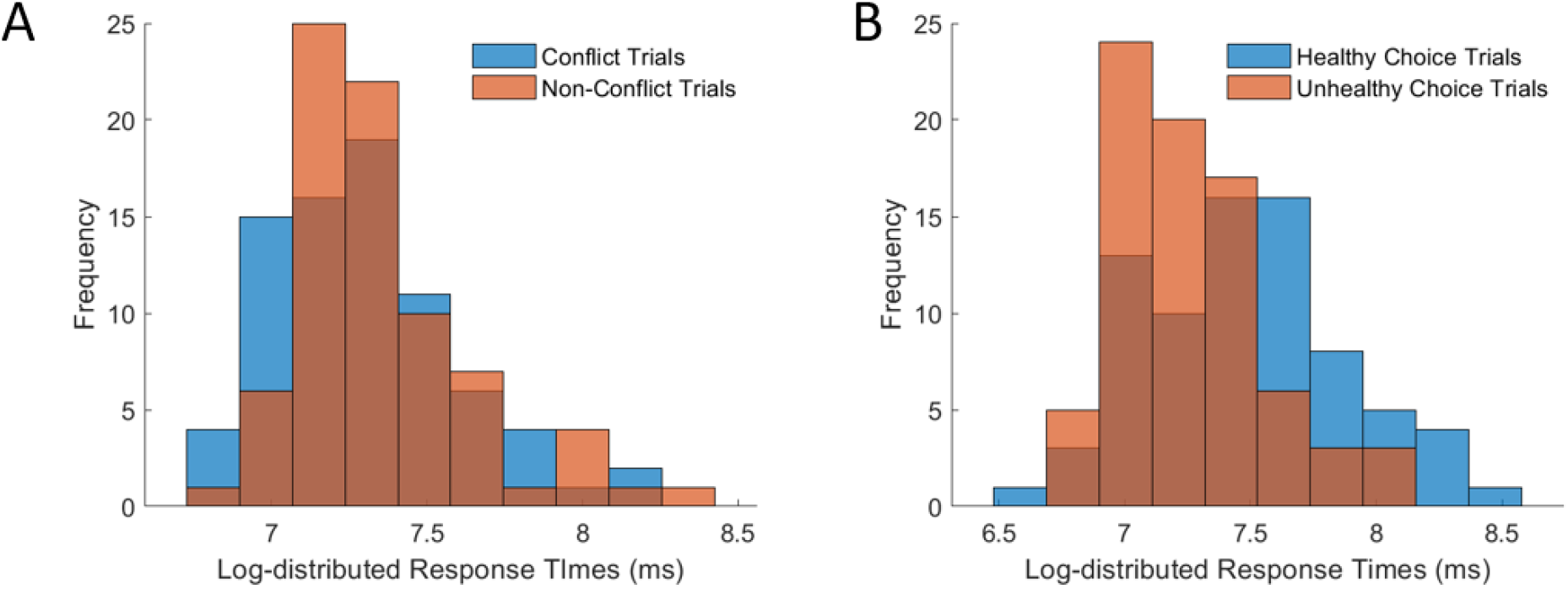
Response time distributions by trial type. A) Log-transformed response time distributions in conflict trials, in which one food was tastier and less healthy than the other, and non-conflict trials. B) Among conflict trials, RT distributions when the healthy or unhealthy choice was made.

**Supplementary Figure 4.**
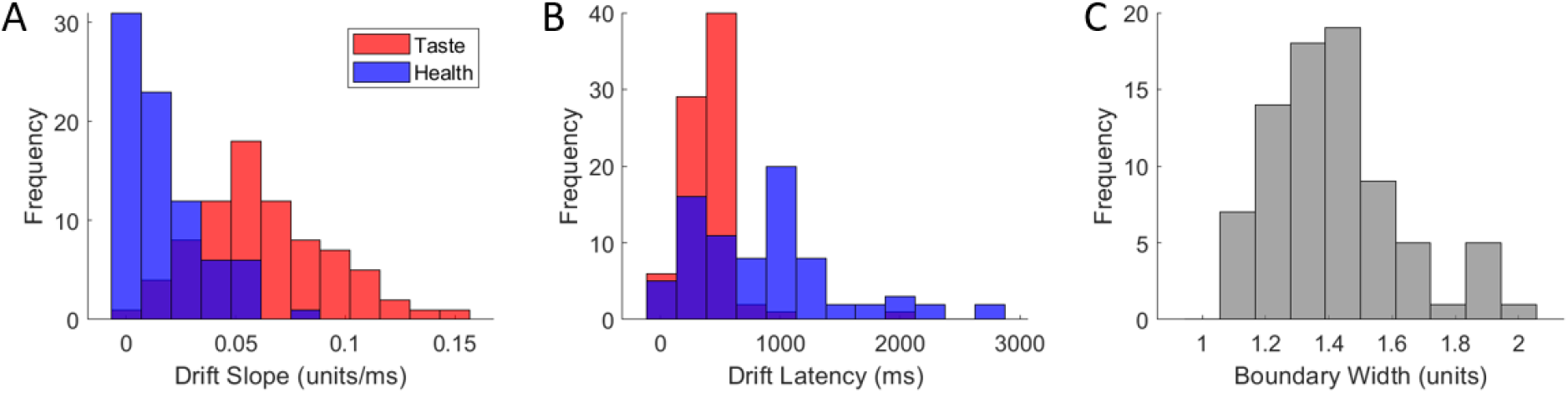
Distribution of best-fitting mtDDM parameters. (**a**) Histogram depicting the distributions of best-fitting taste and health drift slopes (δ_T_ and δ_H_) across participants. (**b**) Distribution of best-fitting taste and health drift latencies (t*_T_ and T*_H_). (**c**) Distribution of estimated boundary size.

**Supplementary Figure 5.**
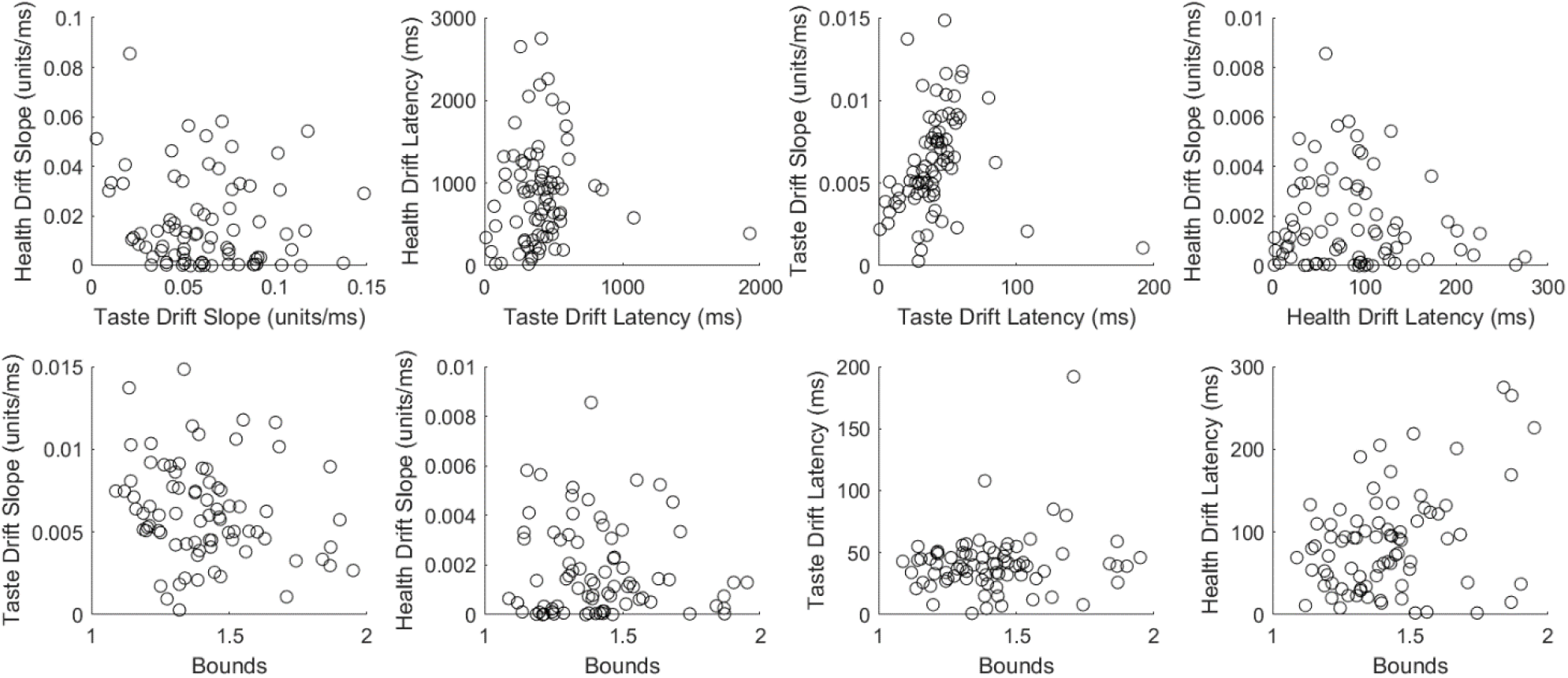
Scatter plots of each free mtDDM parameter combination. Each fitted mtDDM parameter is plotted against each other parameter to visualize the correlation (or lack thereof) between parameters.

**Supplementary Figure 6.**
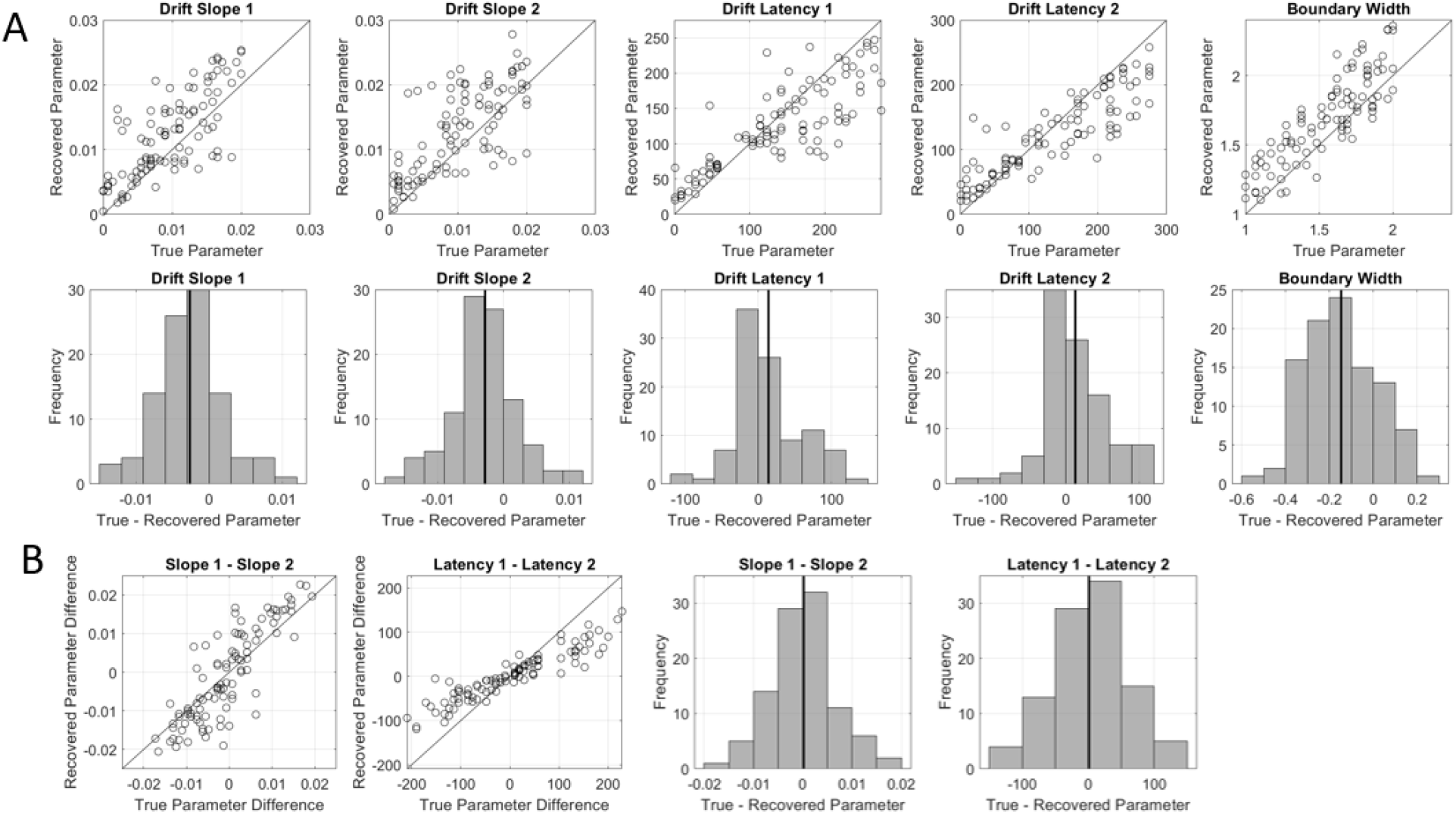
Difference between recovered and true simulated parameters. (**a**) Each “true” mtDDM parameter is plotted against its recovered estimate. The distribution of differences between true and recovered parameters are shown below each scatterplot. (**b**) The difference in Drift Slopes and Latencies for Taste and Health are plotted, with the true parameters of the simulations plotted against their recovered parameters. The distribution of differences in true and recovered parameter differences (Taste-Health) are shown below each scatterplot. In each scatter plot, the black line represents a perfect correlation line. In each histogram, the black line represents the mean difference between true and recovered parameter.

**Supplementary Figure 7.**
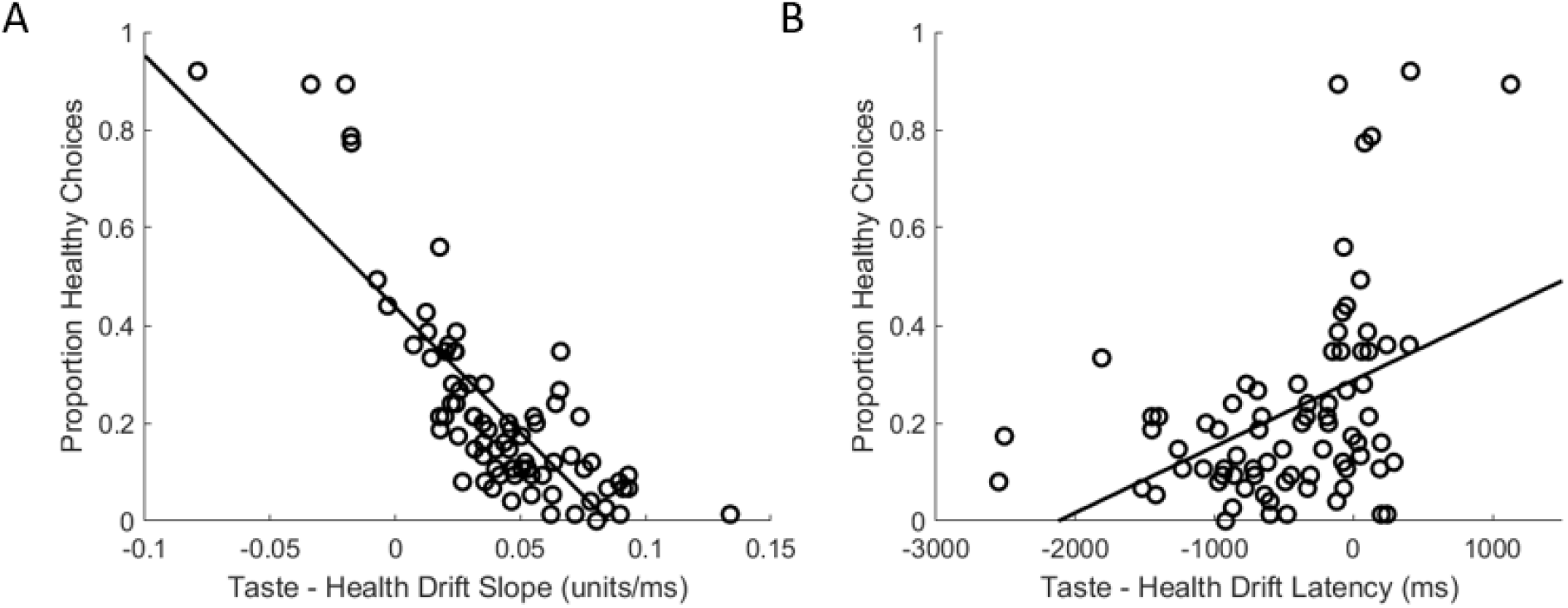
Attribute drift slopes related to choice. (a) Drift slopes plotted as a function of the proportion of healthy choices made. (b) Relative drift latency (taste – health) plotted against proportion healthy choices.

**Supplementary Figure 8.**
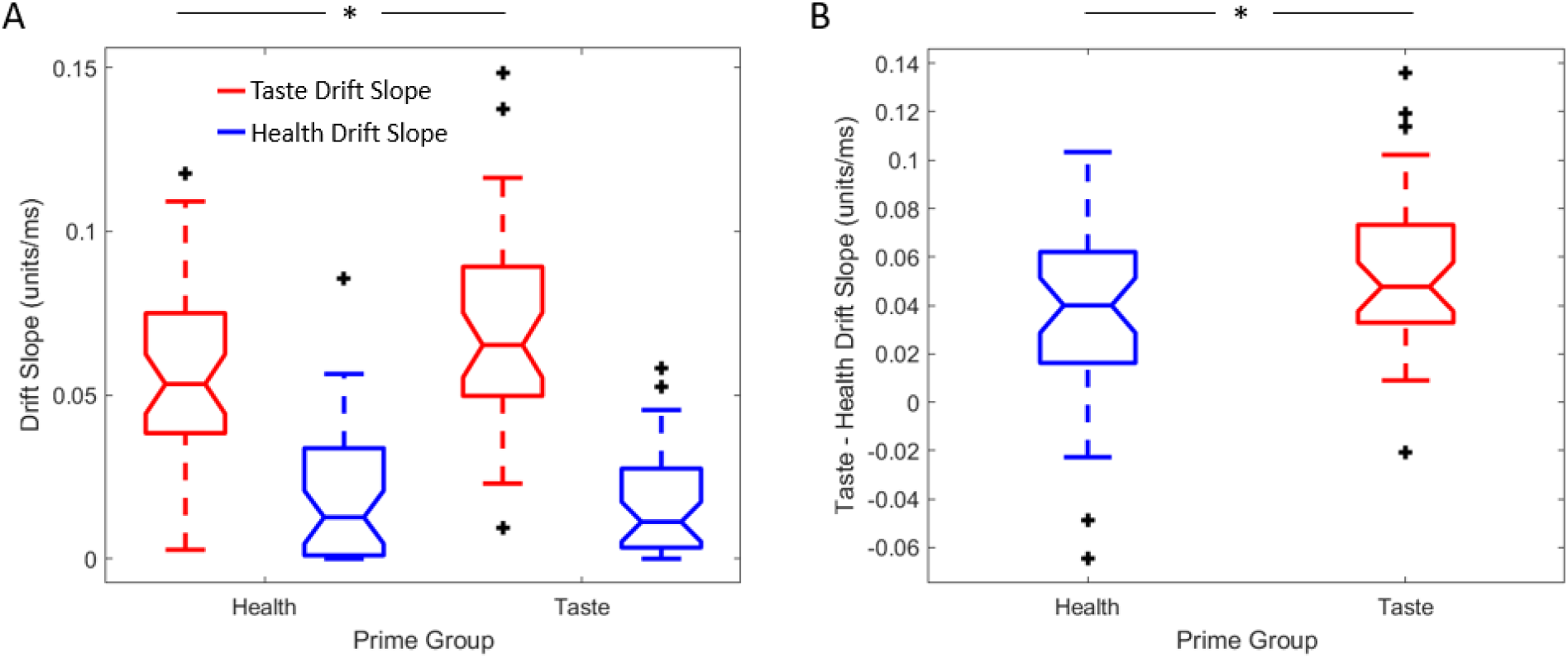
Influence of prime on mtDDM parameters. (**a**) Drift slopes for food tastiness and healthfulness by prime condition. (**b**) Difference in taste and health drift slopes by prime condition. For both plots, black crosses represent outliers as determined by Matlab’s boxplot.

**Supplementary Figure 9.**
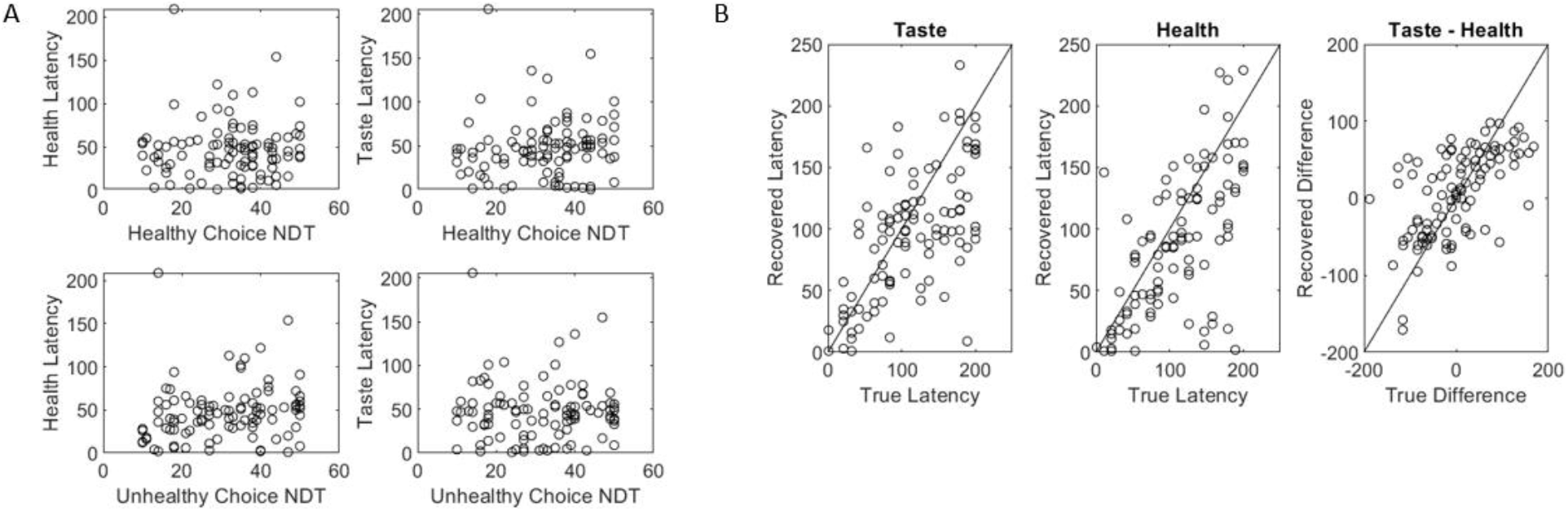
Simulation results from test of bias caused by choice-specific non-decision times (csNDTs). (**a**) Correlation between csNDTs and estimated latencies. (**b**) Correlation between recovered and true latencies when csNDTs are present in the decision process.

## Supplementary Tables

**Table S1.** Model comparisons.

|  | Mean BIC | Std. Dev. BIC | Sum BIC | Min. BIC | Max. BIC |
| --- | --- | --- | --- | --- | --- |
| **mtDDM** | 1111.21 | 97.39 | 87785.52 | 907.64 | 1347.02 |
| **mDDM** | 1143.37 | 99.50 | 90326.29 | 930.55 | 1391.60 |
| **latDDM** | 1166.25 | 95.36 | 92133.80 | 951.86 | 1407.24 |
| **stDDM** | 1644.45 | 115.59 | 129911.40 | 1381.28 | 2005.32 |
| **sDDM** | 1245.58 | 88.38 | 98400.87 | 1081.39 | 1337.95 |

**Table S2.** Association between health’s drift latency, response times, and healthy choices. Models 3 and 4 use latency fit to one half of trials to predict healthy choice on the other half of trials.

|  | Model (1) | Model (2) | Model (3) | Model (4) |
| --- | --- | --- | --- | --- |
| **log(RT)** | $\beta=0.62$<br>[0.49 0.76]<br>$p<0.001$ | $\beta= 0.38$<br>[0.23 0.52]<br>$p<0.001$ | $\beta= 0.39$<br>[0.24 0.54]<br>$p<0.001$ | $\beta=0.13$<br>[-0.11 0.36]<br>$p=0.29$ |
| **Reported Wanting**<br>**(Healthy – Tasty Option)** | | $\beta= 0.92$<br>[0.78 1.07]<br>$p<0.001$ | $\beta=0.92$<br>[0.78 1.06]<br>$p<0.001$ | $\beta=0.92$<br>[0.77 1.06]<br>$p<0.001$ |
| **Health Latency ($t^*_H$)** | | | $\beta= -0.006$<br>[-0.01 -0.002]<br>$p=0.006$ | $\beta=-0.03$<br>[-0.05 -0.013]<br>$p<0.001$ |
| **$t^*_H$ X log(RT)** | | | | $\beta=0.004$<br>[0.001 0.006]<br>$p=0.006$ |
| **Constant** | $\beta=-5.99$<br>[-7.04 -4.95]<br>$p<0.001$ | $\beta= -3.39$<br>[-4.49 -2.30]<br>$p<0.001$ | $\beta=-2.98$<br>[-4.10 -1.85]<br>$p<0.001$ | $\beta=-1.01$<br>[-2.80 0.79]<br>$p=0.27$ |
| **R<sup>2</sup>** | 0.23 | 0.48 | 0.48 | 0.46 |
| **R<sup>2</sup><sub>adj</sub>** | 0.23 | 0.48 | 0.48 | 0.46 |

95% CI in brackets

**Table S3.** Coefficients of a regression predicting healthy choices by stDDM parameters.

|  | **stDDM** |
| --- | --- |
| Taste drift slope | $\beta=-7.28$<br>[-8.75 -5.80]<br>$p<0.001$ |
| Health drift slope | $\beta=4.58$<br>[2.91 6.26]<br>$p<0.001$ |
| Taste stopping time | $\beta=7 \times 10^{-6}$<br>[$2 \times 10^{-5}$ $1 \times 10^{-5}$]<br>$p=0.41$ |
| Health stopping time | $\beta=3 \times 10^{-5}$<br>[$1 \times 10^{-5}$ $4 \times 10^{-5}$]<br>$p<0.001$ |
| Boundary width | $\beta=-0.05$<br>[-0.18 0.07]<br>$p=0.38$ |
| Constant | $\beta=0.52$<br>[0.34 0.71]<br>$p<0.001$ |
| $R^2$ | 0.76 |
| $R^2_{adj}$ | 0.75 |
| 95% CI in brackets |  |

